# Deciphering the mechanical code of genome and epigenome

**DOI:** 10.1101/2020.08.22.262352

**Authors:** Aakash Basu, Dmitriy G. Bobrovnikov, Basilio Cieza, Zan Qureshi, Taekjip Ha

**Affiliations:** Department of Biophysics and Biophysical Chemistry, Johns Hopkins University School of Medicine, Baltimore, MD 21205, USA; Department of Biophysics, Johns Hopkins University, Baltimore, MD 21218, USA; Department of Biomedical Engineering, Johns Hopkins University, Baltimore, MD 21205; Howard Hughes Medical Institute, Baltimore, MD 21205, USA

## Abstract

Sequence features have long been known to influence the local mechanical properties and shapes of DNA. However, a mechanical code (i.e. a comprehensive mapping between DNA sequence and mechanical properties), if it exists, has been difficult to experimentally determine because direct means of measuring the mechanical properties of DNA are typically limited in throughput. Here we use Loop-seq – a recently developed technique to measure the intrinsic cyclizabilities (a proxy for bendability) of DNA fragments in genomic-scale throughput – to characterize the mechanical code. We tabulate how DNA sequence features (distribution patterns of all possible dinucleotides and dinucleotide pairs) influence intrinsic cyclizability, and build a linear model to predict intrinsic cyclizability from sequence. Using our model, we predict that DNA mechanical landscape shapes nucleosome organization around the promoters of various organisms and at the binding site of the transcription factor CTCF, and that hyperperiodic DNA in *C. elegans* leads to globally curved DNA segments. By performing loop-seq on random libraries in the presence or absence of CpG methylation, we show that CpG methylation leads to global stiffening of DNA in a wide sequence context, and predict based on our model that CpG methylation widely changes the mechanical landscape around mouse promoters. It suggests how epigenetic modifications of DNA might alter gene expression and mediate cellular adaptation by affecting critical processes around promoters that require mechanical deformations of DNA, such as nucleosome organization and transcription initiation. Finally, we show that the genetic code and the mechanical code are linked: sequence-dependent mechanical properties of coding DNA constrains the amino acid sequence despite the degeneracy in the genetic code. Our measurements explain why the pattern of nucleosome organization along genes influences the distribution of amino acids in the translated polypeptide.

## Introduction

Two broad strategies have been employed to understand how DNA sequence modulates the bendability and shape of DNA molecules. The first involves compilation of DNA motifs found to be associated with DNA segments of specific curvatures and global shapes, as observed in crystal structures or via other methods. For instance, long polyA tracts occur in straight DNA segments^1^, whereas periodic stretches of poly-As give rise to an average curved configuration^2,3^. Periodic occurrences of dinucleotides at the helical repeat has been shown to characterize highly bent DNA^4^. Along the known nucleosome positioning sequences, TA dinucleotides particularly occurs at locations where the minor groove bends inwards^5^ and where DNA bending is thought to be most challenging^6^, while G/C elements occur at the opposite locations where the major groove faces inwards^7,8^. However, contradictory configurations have also been noted, especially involving GC rich elements at the minor-groove inward locations^5^. Periodic T-A runs in the mu phase genome flanking the strong gyrase binding site have been implicated in facilitating the extensive DNA bending which gyrase action requires^9^, and hyperperiodic DNA in the Ω4 region of the *C. elegans* genome has been structurally observed to be highly curved^10^.

The second strategy involves mechanically manipulating or constraining specific DNA sequences and measuring parameters that describe the response of the DNA molecule. Sensitive methods such as optical and magnetic tweezing^11–13^ have been used to characterize the sequence-averaged properties of DNA such as persistence length, torsional rigidity, structural transitions, etc. Other experimental methods include measuring cyclization rates of short DNA molecules^14^, DNAse I accessibility of DNA minicircles^15,16^, DNA stretching^17^ via AFM or hydrodynamic drag, etc. However, most experimental techniques developed have limited throughput, and have thus been primarily used to study the mechanics of a limited set of functionally relevant DNA sequences such as promoter sequences^18,19^, nucleosomal sequences^20–22^, sequences involved in transcription activation^23^, etc.

Either of the two approaches – compilation of DNA motifs in available structures of DNA in various complexes, and direct low-throughput manipulation experiments – has been used to quantify the extent to which sequence motifs such as dinucleotide steps influence its mechanical properties. The most recent compilation of dinucleotide parameters from available structural data^6^ (twist/ roll/ tilt) has been used to predict plectoneme density along genomic DNA^24^, and similar compilations in the past have been used to build models for DNA mechanics to explain nucleosome organization^25^. Limited throughput DNA cyclization experiments has also been used to assign such parameters and construct a model for the sequence-dependence of DNA bendability^14^.

Computational methods such as Monte Carlo (MC) or Molecular Dynamics (MD) simulations have widely been employed to understand how DNA sequence influences DNA mechanics and shape. All atom MC simulations have been used to predict DNA shape parameters such as tilt, roll, twist, etc, from sequence^26,27^, while MD simulations have observed surprising polymorphic behaviors associated with the shape parameters describing B-DNA^28,29^. However, experimental validation of such predictions is typically limited in sequence context.

Finally, high-throughput SELEX-based methods have been used to identify sequence features that are more enriched or de-enriched in highly loopable molecules by performing several rounds of selection and enrichment of highly cyclizable molecules from an initial random pool of unknown sequences^4,30^. These methods indicate that A/T containing dinucleotides are enriched and CG dinucletides are de-enriched in loopable molecules, and that periodicities in certain dinucleotides such as ApA is a signature of loopable molecules. However, quantitatively measurements of DNA bendability of sequences has not been performed using the method, and consequently quantitative models have not been developed on the basis of this data.

We recently reported the a high-throughput experimental method called loop-seq to determine the cyclization propensity of as many as ∼90,000 different specified 50 bp DNA sequences at a time^31^. DNA fragments immobilized on a surface via biotin-streptavidin linkage are permitted to briefly cyclize *in situ*. This is followed by the selective digestion of unlooped molecules, as performed in SELEX based methods^4^. Molecules that are more cyclizable are thus able to survive this digestion step with higher probability. Via deep sequencing, we calculate the number of surviving copies of each sequence and compare the distribution to that obtained in a control experiment where only the DNA digestion was omitted to obtain the cyclizability of every sequence. The location of the biotin tether modulates the value of cyclizability. To correct for this, cyclizabilities are measured for various tether locations, and the values so obtained are combined using a sinusoidal model to obtain intrinsic cyclizability – a quantity inherent to DNA sequence and independent of external factors. We noted that both dynamic flexibility and static curvatures can contribute to intrinsic cyclizability. Using loop-seq, we reported intrinsic cyclizabilities of DNA fragments of random sequences, as well as fragments that span substantial genomic regions such as those spanning the entire chromosome V and regions around the Transcription Start Sites of hundreds of genes in *S. Cerevisiae*.

Here we apply loop-seq to a large library of randomly specified sequences and correlate the measurements of intrinsic cyclizability with sequence features. We measure “bendability quotients” for all di, tri, and tetranucleotide steps, and further characterize how the pairwise distributions of dinucleotides influence DNA bendability. We also characterize how intrinsic cyclizability is influenced by DNA shape and by cytosine methylation in the CpG context and find that CpG methylation in a broad sequence context makes DNA more rigid. We use this characterization to develop models to predict intrinsic cyclizability from sequence and CpG methylation state, using both a linear and a machine-learning approach. Our models predict a pattern of intrinsic cyclizability around Transcription Start Sites (TSSs) of various organisms that broadly retain universal features that we found earlier to impact nucleosome positioning and remodeling^31^: local peaks in intrinsic cyclizability correlate with the canonical positions of TSS-proxial nucleosomes, and in most cases, a sharp dip in intrinsic cyclizability marks the region just upstream of the TSS. We found that in mouse genes, where CpG sites are relatively more abundant around promoters, CpG methylation leads to a dramatic alteration of the promoter mechanical landscape. It suggests how methylations may influence gene expression by modulating DNA mechanical, and consequently processes like nucleosome organization. We found that highly flexible DNA at CTCF binding sites and in the surrounding region is correlated with nucleosome occupancy, suggesting that sequence-encoded DNA mechanics aids in nucleosome positioning at CTCF sites. We found that highly curved DNA in the Ω4 region of the C. elegans genome is correlated with sequence features we identified as indicative of curved DNA. Finally, we found both on the basis of our predictive model and direct measurements, that while the degeneracy of the genetic code offers some degree of independent control of mechanical and genetic information, the two are linked, leading to correlations between amino acid distribution along a proteins and nucleosome positioning in its coding DNA.

## Results

Loop-seq provides the unprecedented opportunity to use direct, high-throughput, and quantitative experimental data in building a physical understanding of how sequence features or chemical modifications influence DNA mechanics. By performing loop-seq, we previously reported^31^ the intrinsic cyclizabilities of 12,472 50 bp DNA fragments (this set of randomly chosen sequences was termed the ‘random library’ (supplementary note 1)). We now also measured intrinsic cyclizability of all sequences in the random library after we enzymatically methylated all CpG dinucleotides for a side by side comparison. We find that overall, DNA sequences become more rigid upon CpG methylation, thereby implicating CpG methylation as a possible dynamic means of regulating local DNA mechanics, in a wide sequence context (Fig. 1a, Supplementary Note 23). We therefore use these two datasets – intrinsic cyclizability measurements of sequences in the random library, and in the CpG methylated random library, as a starting point for understanding how DNA sequence and epigenetic modifications influence its mechanical properties.

**Figure 1:**
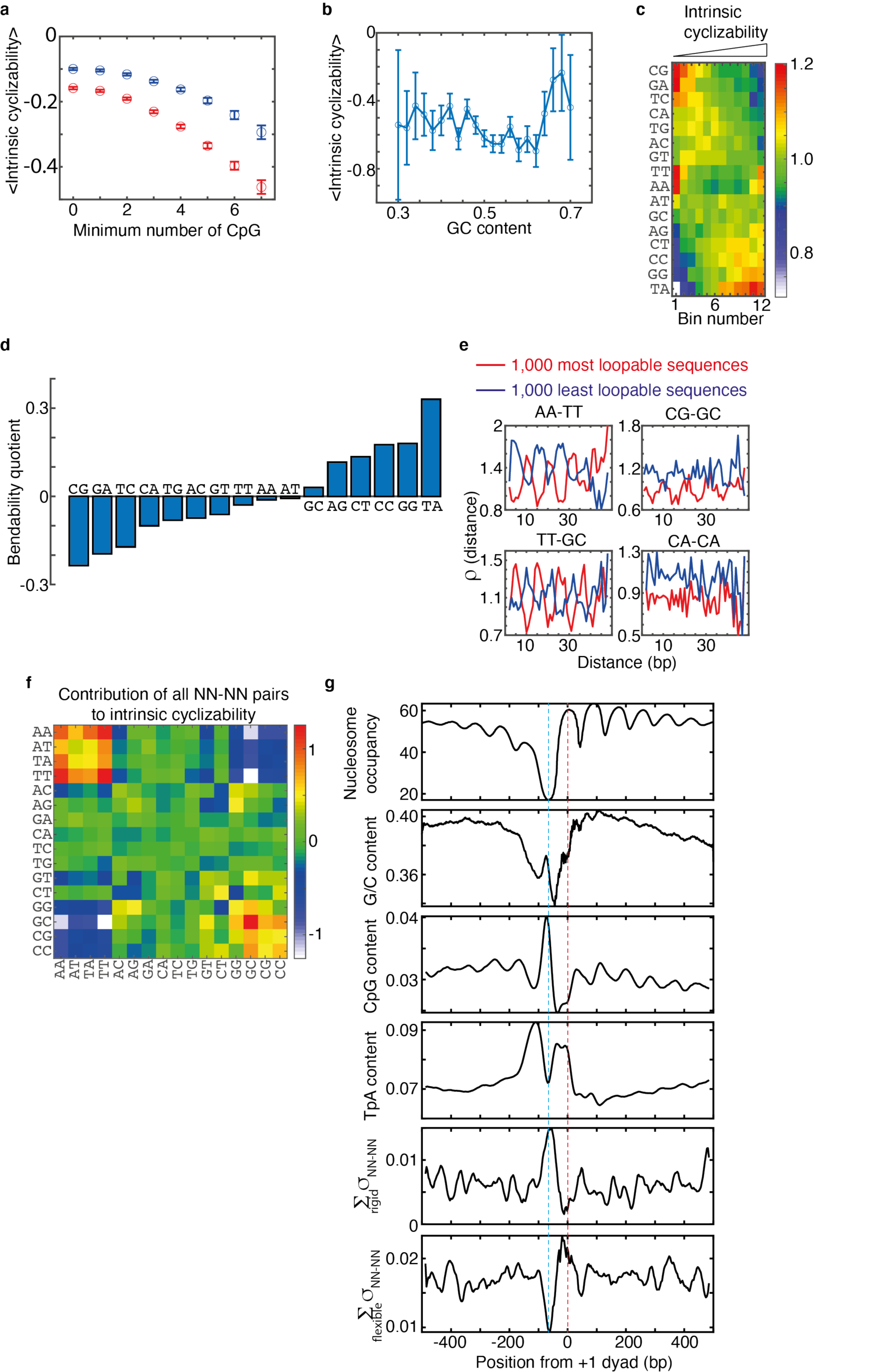
(a) Abscissa are mean intrinsic cyclizabilities of sequences in the random library (blue) and in the methylated random library (red) that have at least the number of CpGs as specified by the ordinate. Intrinsic cyclizability values of the methylated random library have been adjusted to allow for comparison with those of the random library (Supplementary Note 22) (b) Mean intrinsic cyclizability of sequences in the random library as a function of G/C content. Mean is calculated only over those sequences in the random library whose G/C content is as specified by the ordinate. Error bars are s.e.m. (c) The 12,472 sequences in the random library were sorted according to increasing intrinsic cyclizability and grouped into 12 bins with 1,039 sequences each. 4 remaining sequences were ignored. Within each bin, the normalized number of times each of the 16 dinucleotides occur is color coded and depicted (see supplementary note 2). (d) Bendability quotient for a dinucleotide is defined as the slope of the linear fit to the plot of the normalized number of times it occurs among sequences in each of the 12 bins in panel c, vs the mean intrinsic cyclizability of sequences in the 12 bins (supplementary note 2) (e) Pairwise distance distribution functions vs separation distance, averaged over the 1,000 sequences in the random library that have the most (red) or least (blue) values of intrinsic cyclizability, for four different NN-NN pairs. See Supplementary Note 4 for plotting details. (f) For a given NN-NN dinucleotide pair, the best fit linear relationship between its helical separation extent in a sequence and the intrinsic cyclizability of that sequence, for all sequences in the random library, was obtained. The heatmap here depicts the slopes of these linear relationships for all 136 NN-NN pairs. See Supplementary Note 5 for details. (g) Nucleosome occupancy and various sequence parameters as functions of position from the dyad of the +1 nucleosome, averaged over all identified 4,904 genes in *S. cerevisiae* (see supplementary note 6 for plotting details). To calculate ∑_*rigid*_ *σ*_*NN*–*NN*_, the ten NN-NN pairs that make the most negative contribution to intrinsic cyclizability were identified (supplementary note 6). The sum of the helical separation extents of these pairs over a 50 bp DNA fragment centered around the ordinate value was calculated for each gene. The values were averaged to obtain the abscissa value. ∑ _*flexible*_ *σ*_*NN*–*NN*_ was similarly calculated. See supplementary note 6 for heatmaps of TpA and CpG contents.

### Sequence features that influence DNA cyclizability

We found that the simplest sequence feature – overall G/C content – is uncorrelated with intrinsic cyclizability among sequences in the random library, except for an increase at very high G/C content above 65% (Fig. 1b). We next analyzed if and how the total contents of various dinucleotides in a sequence influences its intrinsic cyclizability, ignoring for now how the dinucleotides are distributed along the sequence (Fig 1c). We sorted the 12,472 sequences in the random library according to increasing intrinsic cyclizability, and divided the sorted sequences into 12 bins, each containing ∼1,039 sequences. For a given dinucleotide, we calculated the total number of times it occurs in the 1,039 50 bp sequences in each bin and appropriately normalized these numbers (Supplementary Note 2) (Fig. 1c). As most dinucleotide contents vary monotonically with bin number, we found the best fit linear relationship between the normalized dinucleotide content of a bin (for a particular dinucleotide) and the mean intrinsic cyclizability of all sequences in the bin. We defined its slope as the bendability quotient of that dinucleotide (Fig. 1d, Supplementary Note 2). We find that TpA has the highest bendability quotient, implying that, on average, sequences with high intrinsic cyclizability tend to have more TA steps. CpG has the lowest bendability quotient.

Several earlier works based on low-throughput experiments have suggested that TA dinucleotides are particularly flexible^14,32–35^. Also, consistent with this, TA dinucleotide steps have long been known to be associated with highly bent DNA in nucleosomes^36,37^. Less has been known about the contribution of CG dinucleotides to DNA mechanics. A SELEX-based study found that CG dinucleotides are depleted among the highly loopable sequences, but they did not find specific enrichment of TA dinucleotides over other T/A dinucleotides^4^.

Certain dinucleotides such as TT and AA are represented highly in both the bins of extremely high and the bins of extremely low intrinsic cyclizability (Fig. 1c), suggesting that their distribution along the DNA segment could play a greater role in determining their contribution to intrinsic cyclizability than their overall content alone. Short stretches of dA:dTs bend DNA towards the minor groove. These bends add in or out of phase if repeated at the helical or half-helical period, respectively. In turn, it results in globally curved or straight molecules that would have high or low intrinsic cyclizability, respectively^38–40^.

The availability of high-throughput data from loop-seq permits, for the first time, the tabulation of how phased repeats of a large number of sequence motifs (not just short dA:dT stretches) quantitatively influence DNA mechanics. There are 136 NN-NN dinucleotide pairs (such as AA-TT, AC-GG, etc). For a given pair and a given sequence, we define the pairwise distance distribution function, ρ(*i*), as the number of times that the two dinucleotides in the pair are separated by a distance *i* in the sequence, normalized by an appropriate factor (Supplementary Note 3). We find that the AA-TT dinucleotide pairs have striking oscillations in *ρ(i)* when averaged over thousands of sequences: these dinucleotides tend to more likely be separated by 10, 20 or 30 bp (i.e. multiples of the helical repeat) and less likely be separated by 5, 15 or 25 bp (i.e. odd multiples of half the helical repeat) among the 1,000 most cyclizable sequences (Fig. 1e, red curve). The trend is reversed for the least cyclizable sequences (blue curve). These observations are consistent with the known fact that short dA:dT stretches repeated every 10 bp result in curved molecules. We discovered that other dinucleotide pairs can influence intrinsic cyclizability because of being separated at the helical or half helical repeat (Fig. 1e). The CG - GC pair shows a similar behavior to AA-TT pair. The TT - GC pair show the reverse effects: sequences with high cyclizability have more instances of TT - GC separation at 5, 15, or 25 bp, and less instances at 10, 20, 30 bp, while the reverse is observed for sequences with low intrinsic cyclizability. For several other pairs, such as CA - CA, ρ shows no oscillatory patterns.

The general pattern of curves in figure 1e indicates that NN-NN dinucleotide pairs influence intrinsic cyclizability depending on the degree to which they are separated at even or odd multiples of the half-helical repeat, respectively. To quantify the effects of all possible NN-NN pairs on intrinsic cyclizability, we define a quantity called the helical separation extent of a given NN-NN pair in a given sequence (*σ*_*NN*–*NN*_) as follows:

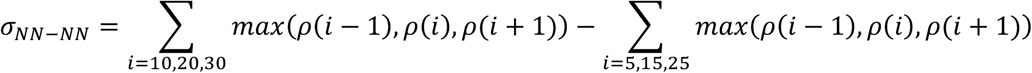

where ρ is the pairwise distance distribution function. If a sequence has a high value of helical separation extent for the NN-NN pair, it implies that the two NNs have high instances of being separated at multiples of the helical repeat, and low instances of being separated at odd multiples of half the helical repeat, in that sequence. For every NN-NN pair, we calculated its helical separation extent among all 12,472 sequences in the random library, found the best-fit linear relationship between helical separation extent and intrinsic cyclizability, and considered its slope as a measure of the contribution that the NN-NN pair makes towards intrinsic cyclizability (Fig. 1f).

We find that NN-NN dinucleotide pairs with all Ns = A or T, make a positive contribution to intrinsic cyclizability (Fig. 1f). In other words, DNA sequences become more or less intrinsically cyclable when such dinucleotides are more often separated by multiples of the helical repeat or odd multiples of half the helical repeat respectively, consistent with earlier reports that short dA:dT stretches at the helical repeat lead to bent DNA. We find a similar effect for some of the cases where all Ns = G or C (fig. 1e) and show that this is not just a reflection of the fact that sequences which have high G/C content every 10 bp also tend to, by exclusion, have high A/T content every 10 bp (supplementary note 20). This novel finding is consistent with an earlier suggestion that G/C dinucleotides tend to bend DNA towards the major groove^14^. These curvatures might similarly add in or out of phase in sequences where G/C rich dinucleotides occur every helical or half helical repeat, respectively. In the case of hybrid pairs, where the one dinucleotide in purely A/T containing and the other purely G/C containing, helical separation extent has an overall negative contribution to intrinsic cyclizability. Consecutive bends towards the minor and major groove every half-helical repeat or helical repeat would add in and out of phase respectively, presumably contributing to this effect. These findings are consistent with aspects of SELEX experiments which demonstrated that A/T rich stretches separated by 5 bp from G/C rich stretches, on average, are more represented in highly loopable molecules^4^. In addition, there are several hybrid pairs, such as AG-GG, or GT-AG, which make significant positive or negative contributions to intrinsic cyclizability, which cannot be described on the basis of any simple expectations.

Both dynamic flexibility as well as static bends may contribute to intrinsic cyclizability as noted previously^31^. Speculatively, the NN-NN helical separation extent contributes to intrinsic cyclizability mainly by altering the static bent structure of DNA and determining whether static bends add in or out of phase. On the other hand, bendability quotients, by virtue of being independent of nucleotide distribution, may reflect how NNs contribute to dynamic flexibility.

We next asked whether the sequence features we have identified as being relevant to intrinsic cycliability are distributed near TSSs of genes in *S. cerevisiae* in a manner consistent with earlier measured intrinsic cyclizability values. Genes are characterized by critical features such as a well-ordered array of downstream nucleosomes (labeled +1, +2, etc, depending on their distance from the TSS) and an upstream Nucleosome Depleted Region (NDR)^41,42^. Loop-seq showed that, averaged over the TSSs of 576 genes in *S*. cerevisiae, genes possess a narrow well-defined region of unusually low intrinsic cyclizability co-centric with the NDR and possess regions of high local intrinsic cyclizabilities at the known canonical locations of the downstream nucleosomes^31^. As nucleosome formation involves extensive DNA bending, our observed pattern of intrinsic cyclizability around TSSs would contribute to nucleosome depletion at the NDR and nucleosome positioning at the canonical locations of gene body nucleosomes. We also showed^31^ that the rigid DNA region co-centric with the NDR aids INO80 in positioning the +1 nucleosome^43,44^.

We identified 4,912 genes in *S. cerevisiae* that have previously been annotated as ORFs and have had both ends mapped with high confidence^45^. Averaged over these genes, we first plot nucleosome occupancy as reported earlier^46^ (fig. 1g), which shows the characteristic dip near the NDR upstream of the +1 nucleosome and the well-ordered set of downstream nucleosomes. We find that overall G/C content shows no special features around the TSSs, other than a general dip near the NDR (Fig. 1g). The dip is likely a result of the fact that NDRs are AT rich. However, the distribution of CG dinucleotides shows a narrow, well-defined peak co-centric with the NDR and local dips at the locations of various nucleosome dyads (Fig. 1g, supplementary note 6). These features cannot be explained on the basis of overall G/C content but are consistent with the CG dinucleotide having the most negative bendability quotient (Fig. 1d). We also find that TA dinucleotide content is high at the locations of the +/-1 nucleosomes and dips near the NDR (Fig. 1g, supplementary note 6), consistent with the TA dinucleotide having the highest bendability quotient (Fig. 1d). We also find that the sum of the helical separation extents of the 10 NN-NN pairs that make the most negative contribution to intrinsic cyclizability (which mostly include hybrid pairs where one NN is A/T containing while the other is G/C containing, Fig. 1f) averaged over all genes in *S. cerevisiae*, sharply peaks near the NDRs and dips near the dyads of the +1 nucleosomes (Fig. 1g). In other words, such pairs tend to more often be separated at the helical repeat (or less often at half the helical repeat) near the NDRs of genes, and the other way around near the dyads of +1 nucleosomes. We can now appreciate the physical significance of this distribution pattern: it contributes to making DNA more straight at the NDRs and more cyclizable near the +1 nucleosome dyad. Likewise, the sum of the helical separation extents of the 10 NN-NN pairs that make the most positive contribution to intrinsic cyclizability (such as AA-TT, GC-GC, etc, Fig. 1f) averaged over all genes, sharply peaks at the dyad of +1 nucleosomes and shows a dip near the NDRs (Fig. 1g).

### Intrinsic cyclizability is related to DNA shape

DNA shape plays a fundamental role in determining the extent of protein-DNA interactions^47^ and the binding specificities of transcription factors^48^ and other proteins including nucleosomes^49^. Widely varying sequences can have similar DNA shapes, and DNA shape has been suggested to be under evolutionary selection^50^. DNA shape has been described by various features that define the local geometry of base pairing, such as helical twist, propeller twist, roll, shift, etc^51^. The determination of the sequence-dependence of shape parameters has been made possible largely via MD and MC simulations, and the observations of the structures of DNA in various complexes. Loop-seq, therefore, may provide a high-throughput experimental platform to validate aspects of these predictions.

High propeller twist has been suggested to be an attribute of rigid DNA^52^, though rigidity here was determined from the static shape of DNA observed in a limited set of crystal structures rather than high throughput dynamic measurements. We predicted various shape parameters across all sequences of the random library using a reported method based on MC simulations^27^. We found that the 1,000 sequences in the random library that have the highest values of intrinsic cyclizability have a lower magnitude of propeller twist than the 1,000 sequences with the lowest values of intrinsic cyclizability (Fig. 2a). Further, the difference is most pronounced near the middle of the 50 bp region. Because the ends of the DNA are involved in hybridization to form loops, sequence features that influence the ability of DNA to loop should have their most pronounced effect if located near the middle of the sequences.

**Figure 2:**
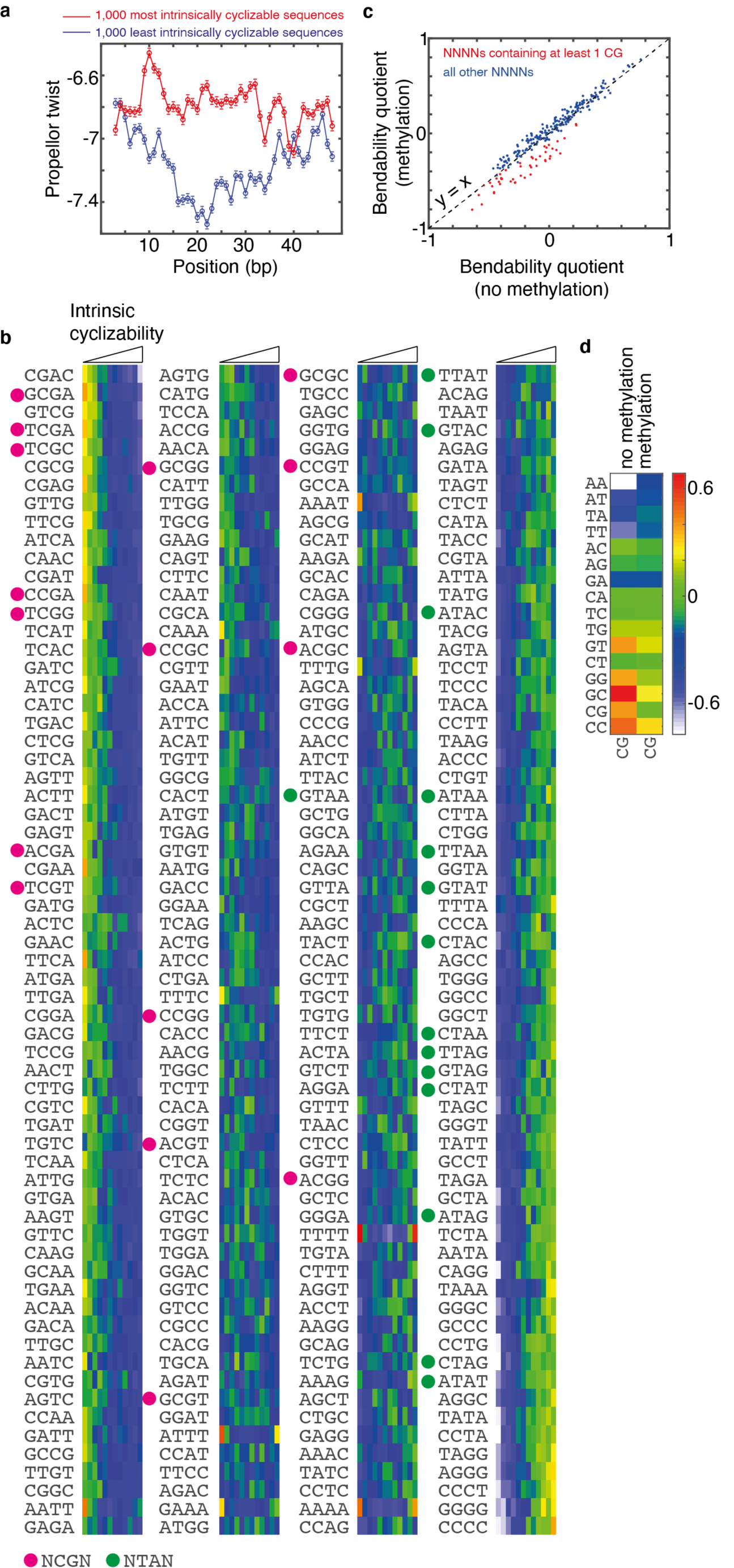
(a) Mean propeller twist as a function of position averaged over those 50 bp DNA fragments in the random library that had the most (red) and least (blue) values of intrinsic cyclizability. Sequences were tiled as a series of pentamers and the associated propeller twist of the central base in the pentamer was assigned on the basis of earlier reports^27^. (b) The 12,472 sequences in the random library were sorted according to increasing intrinsic cyclizability and grouped into 12 bins with 1,039 sequences each. 4 remaining sequences were ignored. Within each bin, the normalized number of times each of the 256 tetranucleotides occur is color coded and depicted (Supplementary Note 7). Tetranucleotides containing a CG in the middle (ie of the form NCGN) are indicated by a magenta circle, while those containing a TA in the middle are indicated by a green circle. (c) Bendability quotients for all 256 NNNNs obtained from the measured values of intrinsic cyclizability of the 12,472 sequences in the random library vs those obtained from the measured values of intrinsic cyclizability of the sequences in the methylated random library (which contain the identical set of 12,472 sequences, except all occurring CpG are cytosine methylated). Dashed line represent x=y. NNNNs are marked in red if at least one CG occurs in it (such as ACGA, CGCG, CGAC, etc). Other NNNNs (such as AAGC, GGGC, etc) are marked in blue. (d) Heatmap representing the contributions of all NN-CG dinucleotide pairs towards intrinsic cyclizability, obtained by considering the intrinsic cyclizability values of sequences in the random library (first column, identical to the 15^th^ column in figure 1f except for the color scale), and obtained from measurements on the methylated random library (second column). Contribution towards intrinsic cyclizability of a NN-CG pair is calculated as done in the case of figure 1f.

### CpG Methylations influences local DNA bendability

The methylation state of CpG dinucleotides are a marker for gene expression and have been shown to also influence the stability of nucleoprotein complexes^53,54^. Recently, low-throughput DNA cyclization measurements and MD simulations suggested that CpG methylation can make DNA more rigid^22^ in a few tested DNA sequences. Apart from the limited throughput, these experiments were also limited in that they did not appreciate the modulatory influence of the biotin tether on DNA looping. Comparisons of intrinsic cyclizability values of sequences in the random vs those in the methylated random libraries showed that CpG methylation, on average, makes DNA more rigid (Fig. 1a). To understand if the specific sequence context the CpG occurs in is important for the overall stiffening effect, we calculated the bendability quotients of all NNNN tetranucleotides in the methylated and the unmethylated libraries. We found that CpG methylation decreases the bendability quotients of all NNNN tetranucleotides that contain at least one CG (Fig. 2c). This finding suggests that CpG methylation results in global stiffening of DNA irrespective of the immediate sequence context it occurs in.

We next looked at the effect of CpG methylation on the contribution of all CG-NN dinucleotide pairs towards intrinsic cyclizability by way of helical separation extent. In the unmethylated case, CG-NN pairs make a positive contribution towards intrinsic cyclizability if N = G or C, and a negative contribution if N = A or T (Fig. 1f). CpG methylation shifts the contribution of the helical separation extent of CG-NN pairs away from extremes, towards zero: CG-NN pairs with G/C rich and A/T rich NNs decrease and increase, respectively, their contribution to intrinsic cyclizability as a result of CpG methylation (Fig. 2d). It is likely that highly positive or negative contribution of dinucleotide pairs towards intrinsic cyclizability arises from consecutive static bending every helical repeat that are either in or out of phase respectively. Therefore, a possible mechanism by which CpG methylation can shift the contribution towards zero is by the suppression of DNA bending towards the major groove at CpG steps. It has indeed been suggested, based on crystal structures, that increased steric hindrance at the major groove as a result of methylation broadens the major groove and suppresses bending^55^.

Loop-seq thus provides evidence that CpG methylation may locally alter the dynamic flexibility and static shapes of DNA molecules. It may therefore serve as a means by which cells can modulate DNA mechanics and thus aid in the altering of biological processes under epigenetic control such as nucleosome organization and gene expression^56^.

### Models to predict intrinsic cyclizability

We have identified various sequence features that influence intrinsic cyclizability, and therefore attempted to build a unified model that takes into account these varied factors to predict intrinsic cyclizability. We first did this by training a shallow neural net with 10 neurons in a single hidden layer (supplementary note 10). DNA sequences were the inputs to the net (Supplementary Note 10). We trained the net based on measurements of intrinsic cyclizabilities of sequences in the Tiling Library, and predicted the intrinsic cyclizability values of sequences in the Random, ChrV, and Cerevisiae Nucleosomal libraries (Fig. 3a). Typical correlation coefficients between predicted and measured values is ∼0.6. Although the neural net model is simple to train and implement, it does not provide any deeper understanding of how various sequence features we identified (Fig. 1) contribute to the final measure of intrinsic cyclizability.

**Figure 3:**
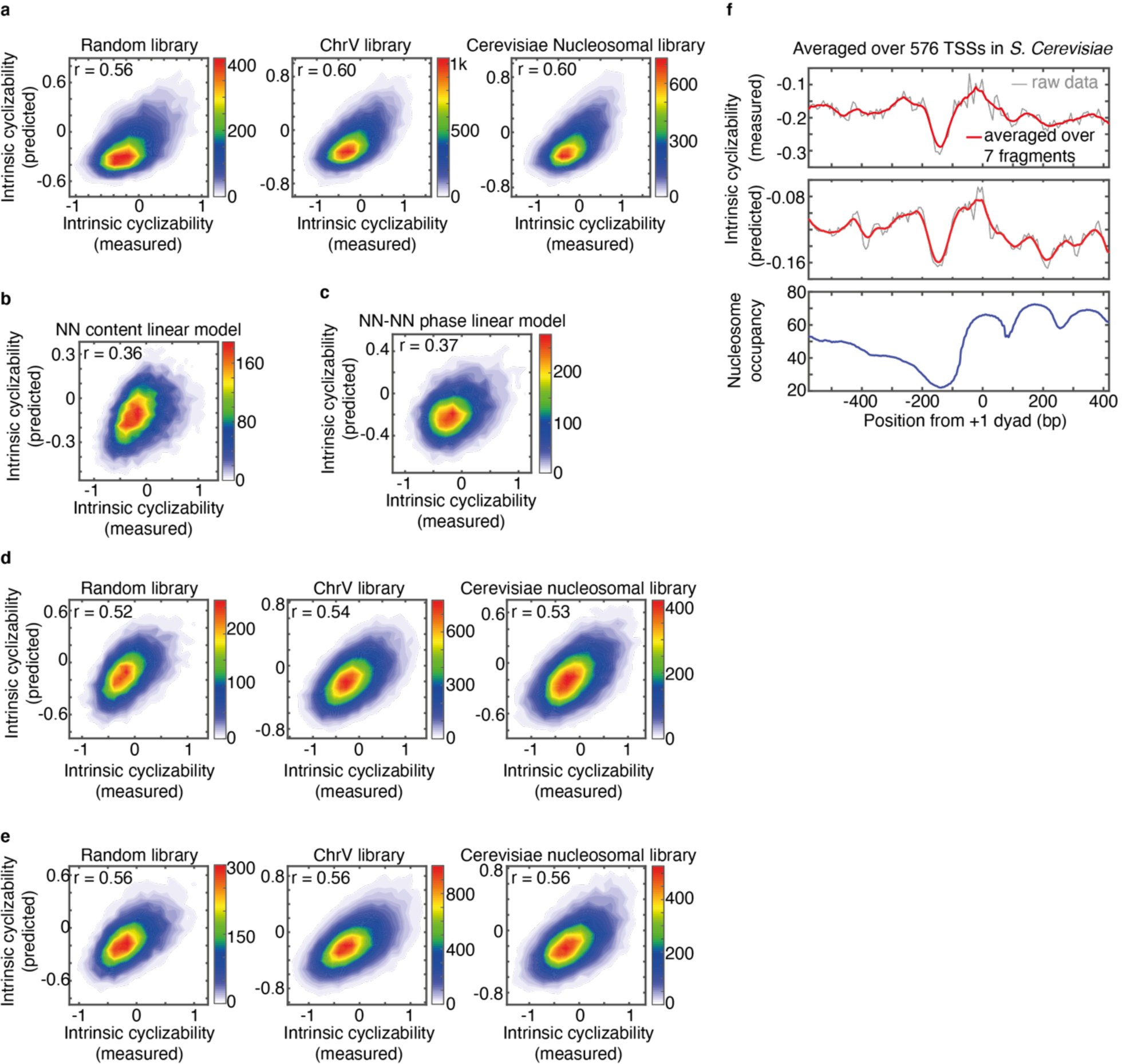
(a) 2-D histograms of scatter plots between measured intrinsic cyclizabilities of sequences in various libraries (Supplementary Note 1) and the associated predicted intrinsic cyclizability. A shallow neural net containing a single hidden layer of 10 neurons was trained using the tiling library to predict intrinsic cyclizability of a 50 bp DNA sequence. Sequences were numerically represented using one hot coding (supplementary note 10) (b) 2-D histogram of the scatter plot between measured intrinsic cyclizabilities of sequences in the random library and the associated predicted intrinsic cyclizability. Here, prediction was made via a model where intrinsic cyclizability of a 50 bp sequence is a linear combination of the number of times each of the 16 dinucleotides occur in the sequence and a constant term (supplementary note 11). Best fit coefficients of the linear model were derived by training the model using the measured intrinsic cyclizability values of sequences in the tiling library (Supplementary Note 1). (c) Same as panel b, except that prediction was made using a linear model where intrinsic cyclizability of a 50 bp sequence is a linear combination of the 136 helical separation extent values in the sequence of the 136 NN-NN pairs, and a constant term (supplementary note 11). (d) 2-D histogram of the scatter plots between measured and predicted intrinsic cyclizability values of sequences in the random, chrV, and cerevisiae nucleosomal libraries (Supplementary Note 1). Here, prediction was made using a model were intrinsic cyclizability of a 50 bp sequence is a linear combination of a constant term and the 16 dinucleotide contents (subject to the constraint that their sum = 49) and 136 helical separation extents that describe the sequence (supplementary note 11). Coefficients were derived by training the model against the tiling library. (e) 2-D histogram of scatter plots between measured and predicted intrinsic cyclizability values of sequences in the Random, ChrV, and Cerevisiae nucleosomal libraries (Supplementary Note 1). Prediction was made using a shallow neural net with a single hidden layer of 10 neurons. 50 bp sequences were represented by a set of 151 parameters (15 of the 16 NN contents, because of the constraint that the sum of all NN contents in a 50 bp sequence must be 49, and the 136 NN-NN helical separation extents) (supplementary note 13). The net was trained using the measured intrinsic cyclizability values of sequences in the tiling library. (f) Measured intrinsic cyclizability, predicted intrinsic cyclizability, and nucleosome occupancy as functions of position from the dyad of the +1 nucleosome, averaged over 576 genes in *S. cerevisiae*. Mean occupancy values^46^ and positions^37^ were as reported earlier. Prediction was performed using the linear physical model trained using the measured intrinsic cyclizability values of the random library. See supplementary note 14 for details.

We therefore attempted to build physical models that are more transparent in how they incorporate the effects of NN content and NN-NN helical separation extent on intrinsic cyclizability (Fig. 1). A 50 bp DNA sequence can be described by a set of 16 NN contents (the number of times each of the 16 dinucleotides occur in the sequence) and 136 NN-NN helical separation extents. We built linear models that treat intrinsic cyclizability as a linear combination of firstly just the 16 NN contents, then just the 136 NN-NN helical separation extents, and finally all the 152 parameters (16 NN contents and 136 NN-NN helical separation extents) (supplementary note 11). The best fit coefficients of this linear combination are obtained by performing multivariant linear regression, using the Tiling Library as the training dataset. We tested the models by predicting the intrinsic cyclizabilities of sequences in the Random, ChrV and Cerevisiae Nucleosomal Libraries (Figs 3b-d). Correlation coefficients between measured and predicted values obtained by the combined linear model are in the range 0.52 – 0.54 (Fig. 3d), close to what was obtained using neural nets (Fig 3a). Finally, we relaxed the condition of linearity by training a shallow neural net with a single hidden layer of 10 neurons to predict intrinsic cyclizability from the 152 input parameters (supplementary note 13) and obtained a small improvement in correlation coefficient to 0.56 (Fig. 3e).

In applying the predictive models to diverse biological contexts, we picked the combined linear model after training it using the measured intrinsic cyclizability values of the Random Library with or without CpG methylation. As a validation, predicted intrinsic cyclizability around 576 genes in *S. cerevisiae* captures all essential features that were earlier identified using the measured intrinsic cyclizability values – a well-defined region of unusually low intrinsic cyclizability co-centric with the NDR, and local peaks in intrinsic cyclizability around the dyads of downstream nucleosomes (Fig. 3f). Although the relative shapes agree, absolute values of intrinsic cyclizability do not agree between prediction and measurement. This is because measured intrinsic cyclizability is defined only up to an additive constant which depends on other sequences present in the library^31^.

### Applications of the predictive model for the sequence-dependence of intrinsic cyclizability

We next applied the combined linear model to predict intrinsic cyclizability around TSSs of genes of four organisms. We predicted intrinsic cyclizability with and without considering CpG methylation and report it together with nucleosome occupancy and overall G/C content around the TSSs of genes in four different organisms (Fig. 4a).

**Figure 4:**
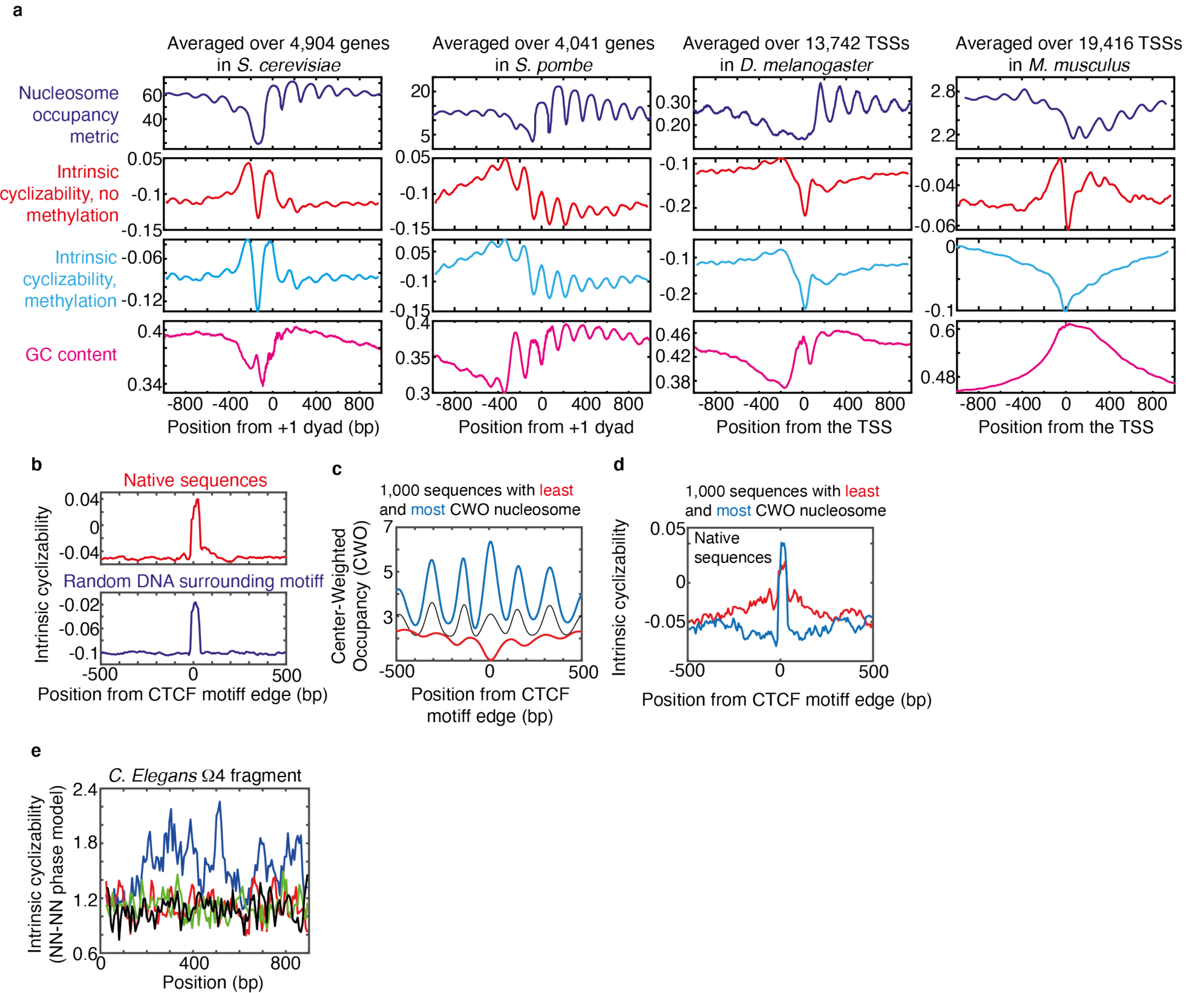
(a) Nucleosome occupancy, predicted intrinsic cyclizability in absence of CpG methylation, predicted intrinsic cyclizability in presence of CpG methylation, and G/C content, as a function of distance from the dyad of the +1 nucleosome (in the case of *S. cerevisiae* and *S. pombe*) or from the TSS (in the case of drosophila and mouse), averaged over a large number of genes in these for organisms. See supplementary note 15 for details. (b) Top panel: Predicted intrinsic cyclizability (predicted by using the linear physical model trained against the random library) as a function of position from the start of the 20 bp CTCF consensus motif, averaged over 19,900 reported CTCF binding sites in mouse embryonic stem cells^63^. Bottom panel: same as the top panel, except DNA outside the 20 bp concensus sequence motif was chosen at random and not obtained from the mouse genome. See Supplementary Note 16 for details. (c) Mean Center Weighted Occupancy (CWO) of nucleosomes as a function of position from the start of the CTCF consensus motif, averaged over all 19,900 CTCF binding sites, and over two sub-groups of 1,000 sites each that have the highest (blue) and lowest (red) CWO value in the region of the CTCF consensus motif. See supplementary note 17 for plotting details. (d) Predicted intrinsic cyclizability as a function of position (where 0 is the start of the CTCF consensus motif), averaged over the two groups of 1,000 sites identified in panel d, that have the highest (blue) and lowest (red) CWO value in the region of the CTCF consensus motif. Plots were obtained in a manner similar to panel b (top). (e) Predicted intrinsic cyclizability and a function of position along the 923 bp Ω4 region of the C. elegans genome^10^ (blue), and along three more suc 923 bp DNA sequences obtained by randomizing the order of nucleotides that occur along the native Ω4 sequence (black, green, red).

In the case of *S. cerevisiae*, predicted intrinsic cyclizability patterns broadly agree with what was measured for the case of 576 genes^31^ (Fig. 3f) – there is a well-defined region of rigid DNA upstream of the +1 dyad, which coincides with the NDR (Fig. 4a). Additionally, clear flexibility peaks are visible at the location of the +1 and -1 nucleosomes, and to a lesser extent, the +2 nucleosome. However, the predicted intrinsic cyclizability map around the +1 nucleosomal dyads^57^ of *S. pombe* is markedly different from that of *S. cerevisiae* in two respects (Fig. 4a). First, there is no special region of rigid DNA upstream of the +1 nucleosome, consistent with some earlier findings that *S. pombe* does not possess a well-defined NDR^57^. Second, intrinsic cyclizability shows very strong oscillations in phase with the nucleosome occupancy oscillation, at least up to +7 nucleosome. Together, these observations suggest that promoter-proximal nucleosomes are positioned mainly through sequence-encoded DNA mechanics of individual nucleosomes instead of through statistical positioning next to an NDR.

Next, we predicted intrinsic cyclizability around TSSs in *D. Melanogaster* and *M. Musculus*, and compared it to the known maps of nucleosome occupancy around TSSs^58,59^. In both cases, a sharply defined region of rigid DNA exists just upstream of the +1 nucleosome, suggesting that a well-defined region of unusually rigid DNA may be a generalizable feature found in many species just upstream of the +1 nucleosome. Further, in drosophila, small peaks in flexibility are also visible at the locations of the +1 and +2 nucleosomes.

We next looked at how CpG methylation would change the mechanical landscape around TSSs in the four organisms. In the cases of *S. cerevisiae, S. pombe*, and *D. melanogaster*, CpG methylation does not cause any significant change in the overall predicted landscape of intrinsic cyclizability around TSSs (Fig. 4a). CpG methylation also has not been reported to occur in these organisms. However, methylation significantly alters the landscape around TSSs in mouse (Fig. 4a): the sharp dip in flexibility near the TSS is no longer prominent. Speculatively, it is possible that the altered DNA mechanical landscape alters nucleosome positioning and the activities of other enzymes, in turn contributing to decreased gene expression.

The transcription factors CTCF is implicated in the maintenance of global chromatin structure and nucleosome organization^60^. The nucleosome remodeler SNF2H associates with CTCF, aiding its binding to the CTCF binding site and the establishment of an ordered phased arrays of nucleosomes around the site^61^. CTCF depletion considerably reduced nucleosome phasing, but increases nucleosome occupancy at the CTCF binding site^61^. Likewise, loss of SNF2H also leads to increase in nucleosome occupancy at the CTCF binding site^62^. Finally, chemical cleavage maps suggest that CTCF binding sites in mouse embryonic stem cells may be occupied by a fragile nucleosome, and sites with stronger CTCF binding signals also have a stronger nucleosomal signal (presumably though not at the same time)^59^ (Fig. 4b). What then leads to high nucleosome occupancy at CTCF binding sites? We hypothesized that sequence-encoded intrinsic cycliability of DNA may inherently favor the formation of a well-positioned and occupied nucleosome at the CTCF sites by facilitating DNA bending, and DNA bendability might also be a requirement for CTCF binding.

To test this hypothesis, we predicted intrinsic cyclizability in the 2,000 bp region surrounding 19,900 identified CTCF consensus motifs in mouse^59,63^. Our model predicts that CTCF binding sites have higher DNA cyclizability than surrounding DNA (Fig. 4b). Further, although the consensus motif of the CTCF binding site extends for 20 bp, DNA up to ∼150 bp away from this region is still, on average, more bendable than surrounding DNA (Fig. 4b), and this is not an artifact of any averaging process (supplementary note 16). The fact that the length of a structured DNA region exceeds the CTCF consensus motif has been recently reported^64^. As nucleosome formation requires extensive DNA bending, our finding suggests that the sequence-encoded property of CTCF binding sites as being highly cyclizable may favor the formation of nucleosomes.

We further tested the idea that DNA mechanics at the location of the CTCF binding site could favor the formation of a well-occupied nucleosome. Nucleosome occupancy (specifically Center Weighted Occupancy (CWO)) has been reported across the genome of mouse embryonic stem cells^59^. We identified two groups of 1,000 CTCF binding sites which have the highest (group 1) and lowest (group 2) nucleosome Center Weighted Occupancy values at the CTCF binding site location, from among the 19,900 CTCF binding sites. Incidentally, group 1 sites also have a much higher degree of nucleosome phasing and ordering on either side of the CTCF motif (Fig. 4c). We asked if predicted intrinsic cyclizability values of DNA around the CTCF binding sites in the two group could explain why group 1 sites possess a better occupied nucleosome. The combined linear model indicates that the CTCF consensus motifs in group 1 sites confer higher local intrinsic cyclizability to DNA than in group 2 sites, and further the contrast in intrinsic cyclizability at the CTCF motif and in the immediate neighboring region is higher for group 1 sites than for group 2 (Fig. 4d). Both of these observations are consistent with group 1 CTCF binding sites having higher nucleosome occupancy than group 2 sites. A well-positioned nucleosome can also be used by nucleosome remodelers, possibly SNF2H, to establish will ordered arrays by evenly spacing nucleosomes on either side of it, as seen in the case of group 1 sites (Fig. 4c). Group 2 sites, however, have overall higher DNA bendability in the region just outside of the 20 bp CTCF consensus motif (Fig. 4d). It may explain why instead of a single well-positioned nucleosome at the CTCF site, group 2 sites have two weaker-positioned nucleosomes straddling it (Fig. 4c). Once again, G/C content alone cannot explain these observations (Supplementary Note 22).

Hyperperiodic DNA in the Ω4 region of the genome of *C. elegans* has been reported to result in globally curved DNA structures^10^. We use the NN-NN phased repeat model to investigate if the presence of phased repeats of dinucleotides favor high intrinsic cyclizability in the Ω4 region. Indeed, the predicted intrinsic cyclizability across the hyperperiodic segment is significantly higher than when the nucleotide order in the ∼900 bp region is scrambled (Fig. 4e).

### DNA mechanics and amino acid sequence are linked

As DNA sequence influences intrinsic cyclizability, we asked to what extent the degeneracy of the genetic code allows for independent control of amino acid sequence and the mechanical properties of coding DNA. We first repeated the evaluation of bendability quotients (Fig. 2b and supplementary note 7) in the context of trinucleotides (which serve as codons) (Fig. 5a). We find that for most amino acids, there is a reasonable amount of spread in bendability quotients among its synonymous codons, suggesting at least a certain degree of independent control of DNA mechanics and protein sequence. However, this is not the case for all amino acids (see for example, Y and D) (Fig. 5a), suggesting that the requirement for DNA in coding regions to have certain values of intrinsic cyclizability might constrain the corresponding amino acid sequence and vice versa.

**Figure 5:**
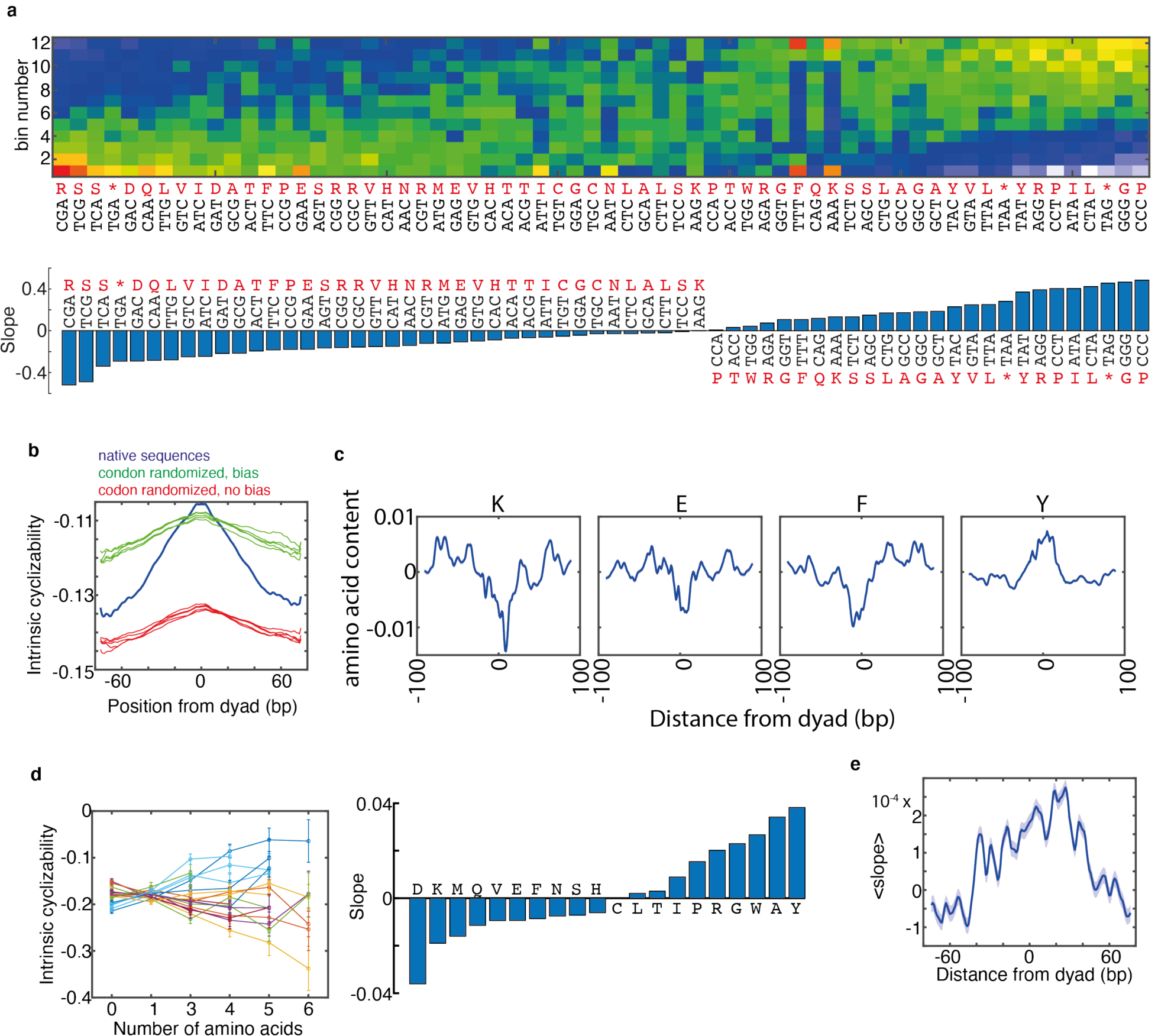
(a) Sequences in the random library were grouped into 12 binds of 1,039 sequences each according to increasing intrinsic cyclizability. Indicated in color is the normalized content of all 64 trinucleotides among the sequences in each of the 12 bins. The analysis is identical to that performed in the context of tetranucleotides in Fig. 2b. Indicated also are amino acids that are coded for by these trinucleotides when they act as codons. Bendability quotients calculated from the normalized contents of each trinucleotide in each bin are depicted in the bar plot and listed in the supplementary file “Bendability_quotients_NNN.txt”. (b) Predicted intrinsic cyclizability in the 200 bp region straddling the dyads of 13,310 nucleosomes in *S. cerevisiae*. These nucleosomes lie along the 4,912 identified genes in *S. cerevisiae* (supplementary note 6), and are classified as +5 through +9 (i.e., these nucleosomes are the 5^th^ through 9^th^ nucleosomes downstream of the TSSs, and lying upstream of the TTSs of these genes) (blue). 10 sets of codon-randomized sequences straddling these 13,310 nucleosomal dyads were also generated, where synonymous codons were chosen while preserving the underlying amino acid sequence. In 5 of these cases, synonymous codons were chosen according to the inherent codon usage frequency in *S. cerevisiae* (green), while in the remaining 5 sets (red), all possible synonymous codons were chosen with equal probabilities. Choosing a codon already present natively was permitted. Intrinsic cyclizabilities averaged across the 13,310 sequences, were predicted for these 10 codon-randomized sets. (c) Contents of various amino acids as functions of distance from the dyads of the 13,310 nucleosomes identified in panel b. (d) Mean intrinsic cyclizability of sequences in the chrV library, as a function of the number of times a specific amino acid occurs in the equivalent in-frame translated amino acid sequences. Mean is calculated over those sequences that lie within coding regions of *S. cerevisiae* chromosome V, and where the number of times the amino acid in question occurs in the equivalent in-frame translated amino acid sequence is as specified by the ordinate. Error bars are s.e.m. Slopes of the straight-line fits to the 20 curves are indicated in the bar plot. (e) Mean amino acid slope along the 13,310 nucleosomes identified in panel b. Slope values are obtained from panel b. See supplementary note 19 for plotting details.

We showed earlier that, for *S. cerevisae*, DNA in the region of deeper gene body nucleosomes (+5 onwards) have a particularly high contrast in intrinsic cyclizability between the dyads and edges, higher in the dyads. We also showed that if the DNA sequences in these regions are altered by randomly choosing different synonymous codons (with or without consideration for the natural codon usage frequency), the contrast disappears, suggesting that the evolution of codon choice has been impacted by the requirement to create a favorable mechanical landscape for the organization of gene body nucleosomes.

We now used the combined linear model to predict intrinsic cyclizability along the nucleosomal regions of all identified +5 - +9 nucleosomes in *S. cerevisiae* (Fig. 5b). We then constructed 10 sets of codon-randomized sequences spanning these nucleosomes. In 5 of these sets, synonymous codons were selected according to probabilities specified by the natural codon usage frequency in *S. cerevisiae*, while in the other sets, they were selected randomly. Codon usage frequency in *S. cerevisiae* leads to a higher overall predicted intrinsic cyclizability than if all synonymous codons were used with equal frequency. In fact, experimental measurements roughly agree with this trend, but high noise in the data obscured a definitive conclusion^31^. This, however, is not generally true for all organisms (supplementary note 21). We next observe that contrast in intrinsic cyclizability between the dyad and the edges is significantly diminished for the codon altered sequences, consistent with experimental observations^31^. However, we find that although contrast is diminished, it is not absent. This suggests that the ability to choose among synonymous codons cannot completely decouple DNA mechanics from the choice of aminoacids coded in gene-body nucleosomes(Fig. 5b). Once again, on the basis of experimental measurements^31^, it was hard to determine if in the codon-randomized sets, the contrast was diminished or totally absent.

Our predictions thus suggest that gene body nucleosomes must have nonuniform distribution of amino-acids across their 147 bp length. To test this, we plotted the average content of all 20 amino acids along the nucleosomal regions of +5 - +9 nucleosomes of all identified genes in *S. cerevisiae*. While most amino acids have no special distribution pattern along nucleosomes (supplementary note), certain amino acids are indeed under or overrepresented at nucleosomal dyads as compared to edges (Fig. 5c).

Our analysis so far suggests that amino acid sequence, to an extent, influences the intrinsic cyclizability of the DNA sequence that codes for it, and the degeneracy in the genetic code cannot completely decouple the two. Previously, we reported intrinsic cyclizability measured along the entire *S. cerevisiae* chromosome V at 7 bp resolution^31^. For selected 50 fragments that lie entirely within coding regions of chromosome V, we determined the associated in-frame amino acid sequence. For every amino acid, we determined the best fit linear relationship between the number of times it occurs in such a sequence and the intrinsic cyclizability of the associated DNA sequence. We treat the slope as a measure of the contribution of that amino acid to intrinsic cyclizability (Fig. 5d). Were the amino acid sequences decoupled from intrinsic cyclizability, most slopes would have been ∼0. We instead find that with increasing amounts of D or Y occurring locally, the measured local DNA intrinsic cyclizability also decreases or increases, respectively.

We next asked whether the amino acid contribution to intrinsic cyclizability has influenced the amino acid distribution in nucleosomal regions of the coding *S. cerevisiae* genome. We aligned the amino acid sequences of all gene body nucleosomes in *S. cerevisiae* to the dyad locations and replaced each amino acid by the slope representing its contribution to intrinsic cyclizability (Fig. 5d). Averaged among all such nucleosomes, we find that the mean slope is higher in regions surrounding the dyad, and lower near the edges of nucleosomes, implying that amino acids that make positive contributions to intrinsic cyclizability (such as Y, Fig. 5c) are more abundant near nucleosomal dyads than edges (Fig. 5e). The slope contributions of every amino acid to intrinsic cyclizability can also be calculated from measurements of intrinsic cyclizability of the sequences in the random library (Suppmentary note 21). Doing so, we still obtained the same result that amino acids which make positive contributions to intrinsic cyclizability are more represented near the dyads of nucleosomes in yeast coding regions.

## Conclusion

Loop-seq provides the unprecedented opportunity to investigate how sequence features and DNA chemical modifications influence the mechanical properties of DNA, on the basis of high-throughput experimental measurements. We identify features such as DNA sequence and shape that may influence DNA mechanics by influencing local DNA bending or dynamic flexibility, and also find that CpG methylation may be a global switch for the modulation of mechanical properties. We show that variations in intrinsic cyclizability are likely to be functionally important in several contexts such as nucleosome organization around TSSs and CTCF sites. Finally, we suggest that the mechanical code is not fully independent of the genetic code. Modern genomic sequences may have been a product of selective pressure shaping both genetic information pertaining to amino acid sequences, and mechanical information which likely regulates the maintenance of, and access to, genetic information.

## Author contributions

AB and TH designed research. AB performed research, analyzed data, and built the predictive models. AB and TH wrote the paper. DGB compiled nucleosome occupancy data from various organisms. BC investigated pairwise correlation among NN-NN dinucleotide pairs in highly loopable and rigid sequences. ZQ assisted with library preparation loop-seq experiments pertaining to the random library.

## Data Availability

All raw data and accession codes will be made available upon request.

## Code availability

All codes used to analyze sequencing data will be made available upon reasonable request.

## Acknowledgements

A.B. and T.H. would like to thank Jun S. Song for suggestions and insights pertaining to developing the linear predictive models. This work was supported by the National Science Foundation Grants PHY-1430124 and EFMA 1933303 (to T.H.) and the National Institutes of Health Grant GM122569 (to T.H.). T.H. is an Investigator with the Howard Hughes Medical Institute.

