## Supplementary notes for "Deciphering the mechanical code of genome and epigenome"

**Supplementary Note 1:**
**Loop-seq and a description of various libraries used**

**<u>The loop-seq protocol:</u>**

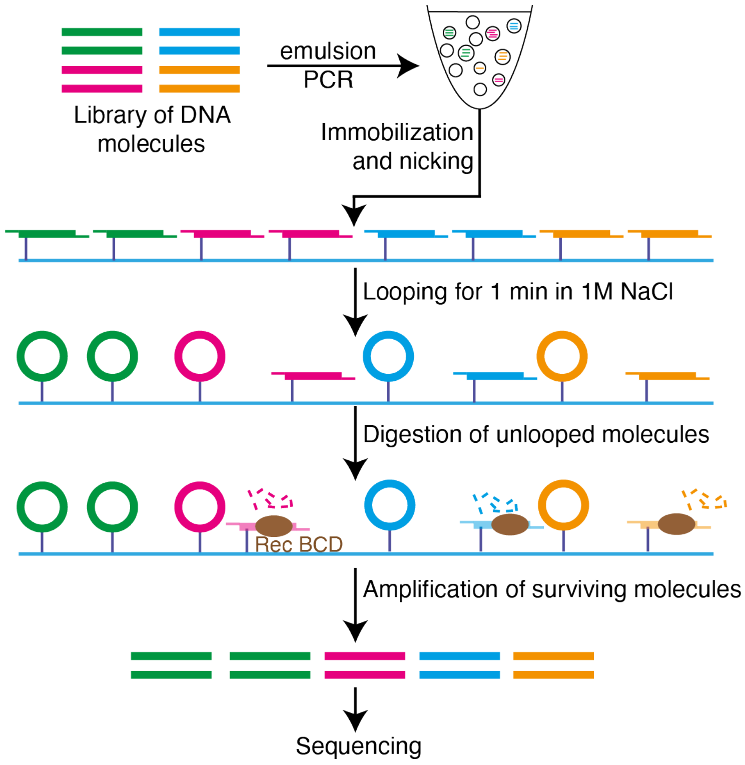


The loop-seq protocol followed in this study is identical to what was described earlier. An initial library of 120 bp DNA fragments, containing up to ~90,000 different DNA sequences is amplified via emulsion PCR. The initial population distribution among the various sequences may be variable. As a demonstration, in this cartoon, an initial library containing 2 copies each of 4 different DNA sequences (in the four colors) is depicted. The central 50 bp are variable among the various sequences in the library, while the flanking 35 bp on either side are identical among all library members. The library was amplified via emulsion PCR and each molecule was chemically modified with a biotin moiety near each end. The library was then surface immobilized on to streptavidin coated beads, and nicked in situ to convert 120 bp duplexes into 100 bp duplexes with 10 nucleotide (nt) flanking complementary single-stranded overhangs, as described. Molecules are permitted to loop for 1 minute in presence of high [NaCl], which permitted stable hybridization of the ends. After a minute, the exonuclease RecBCD is introduced, which digests unlooped molecules while preserving the nicked circular molecules. The library is thus enriched for the more cyclizable sequences. The surviving molecules are amplified and sequenced, and the relative population of each sequence is determined.

In the example shown here, both DNA molecules bearing the "green" sequence survive digestion, while one each of the molecules of other colors do. As a result, even though all sequences had equal relative population in the original pool, the green sequence now has a higher distribution in the selected pool.

As a control, an identically prepared sample was taken through these steps, with the exception that no digestion as performed.

The cyclizability of a sequence was defined as the natural logarithm of the ratio of the relative population of that sequence in the selected pool to that in the control pool. This quantity depends on the cyclizability of other sequences present in the library. For example, in an extreme case of a library containing sequence A plus other sequences all with higher cyclizability than sequence A, sequence A would appear to have an extremely low value of cyclizability even though in a random library sequence A may even be more cyclizable than average. Therefore, cyclizability of a specific sequence measured as part of different libraries would be offset by a constant, and as a result, cyclizability cannot be directly compared between members of different libraries.

We also noted that the distance of the biotin tether from the molecule's end had modulated the measured value of intrinsic cyclizability in a sinusoidal fashion. We corrected for this modulating by measuring cyclizability of a sequence by performing loop-seq three times, each time placing the biotin at a different distance from the ends of all molecules in the library. Using these three measurements, we determined the mean, amplitude, and phase terms associated with this modulation and defined the mean term as the intrinsic cyclizability of the sequence.

Description of various libraries:

All DNA libraries comprised a central 50 bp variable region, flanked by identical adapters for PCR amplification and hybridization during looping.

1. The Random Library

This library comprised 12,472 different DNA sequences. The sequences in the central 50 bp variable regions were specified at random by selection nucleotides with equal probabilities (MATLAB). This library was primarily used to determine how sequence features influence intrinsic cyclizability.

2. The Methylated Random Library

The Random Library was entirely CpG methylated, and loop-seq was performed again to determine intrinsic cyclizability of each sequence.

3. The ChrV library

This library was used to determine how intrinsic cyclizability of DNA varies along the entire length of *S. cerevisiae* chromosome V. The library comprised 82,404 fragments. The central 50 bp variable regions tiled the entire length of *S. cerevisiae* chromosome V at 7 bp resolution:

Sequence 1: position 1 till 50 of chromosome V
Sequence 2: position 8 till 57 of chromosome V
…
Sequence 82,404: position 576,822 till 576,871 of chromosome V

4. The Tiling Library

This library was used to determine how intrinsic cyclizability of DNA varies around the TSSs of genes in *S. cerevisiae*. A list of 576 genes were selected as described before. For every gene, a 2,001 bp region was selected that extends from the location dyad – 1000 till dyad + 1,000, where dyad is the location of the dyad of the +1 nucleosome of that gene (obtained from an earlier report[1] and converted to SacCer3 assembly using liftover[2]).

From this 2,001 bp region, the following 50 bp regions were selected:

Segment 1: position 400 till 449
Segment 2: position 407 till 456
…
Segment 143: position 1,394 till 1,443

This process was repeated for all 576 genes, to obtain a total of 82,368 fragments.

5. The Cerevisiae Nucleosomal Library

This library was constructed to understand quantify the differences in flexibility on either sides of the dyads of well-positioned nucleosomes. The sequences of the 50 bp DNA fragments that lie immediately to the left and to the right of the dyads of the 10,000 nucleosomes in *S. cerevisiae* with the highest reported NCP scores[1] were noted (in the SacCer3 assembly). The dyad itself was not included in these sequences. 19,907 sequences from among 20,000 sequences were selected to comprise this library.

**Supplementary note 2:**
**Obtaining bendability quotients**

Intrinsic cyclizability values of all 12,472 sequences in the random library were sorted and divided into 12 bins of 1,039 sequences each. Bin 1 thus contained the 1,039 sequences with the lowest values of intrinsic cyclizability and bin 12 contained the 1,039 sequences with the highest values of intrinsic cyclizability. As 1039 x 12 = 12,468, the last 4 sequences (in the sorted list of 12,472 sequences) were ignored.

For each of the 16 possible NN dinucleotides, we determined a set of 12 numbers: the number of times it occurs among all the sequences in each bin. As an example, say for the dinucleotide CG, these numbers are $CG_1$, $CG_2$, …, $CG_{12}$.

We then determined, on average, how many times each dinucleotide occurs in a totally random set of 1,039 sequences. To do so, we randomly generated a set of 1,039 x 100 = 103,900 50 bp sequences. Among these sequences, we obtained how many times each of the 16 dinucleotides occur, and divided these 16 numbers each by 100 to determine how many times, on average, each dinucleotide occurs in a random set of 1,039 sequences. As an example, say the CG dinucleotide occurs N times in this random set of 103,900 50 bp sequences. Then N/100 is the mean number of times CG occurs in a random set of 1,039 sequences. The set of numbers $CG_1$, $CG_2$, …, $CG_{12}$ are then normalized by dividing by the quantity N/100.

For each dinucleotide, this normalized set of numbers representing how often they are represented in the sequences within each bin is color coded and displayed in figure 1b.

To determine bendability quotient, for each dinucleotide, we construct a scatter plot of the 12 normalized contents (in each of the 12 bins) vs the 12 mean intrinsic cyclizability values of the 1,039 sequences in each bin. We fit it to a straight line and define its slope as the bendability quotient for that dinucleotide.

The above analysis can be repeated to obtain the normalized contents of all NNN trinucleotides or all NNNN tetranucleotides along the bins. Likewise, bendability quotients associated with tri and tetranucleotides can also be calculated.

#### Supplementary Note 3:
#### Calculation of ρ(i)

$\rho(i)$, for $i$ in the range [1 bp, 48 bp] for each of the 12,478 50 bp sequences in the random library (48 bp is the maximum, non-overlapping, separation distance between two dinucleotides in a 50 bp sequence) is obtained as follows: Say we want to calculate $\rho(i)$ for a dinucleotide pair $N_1N_2$-$N_3N_4$ (this could, for instance, be AC-TT) in a given 50 bp sequence. We generate two row matrices, vec1 and vec2, with 49 columns each, and populated entirely with zeros. Wherever along the 50 bp sequence the dinucleotide $N_1N_2$ occurs, we replace the 0 with 1 in vec1. Similarly, we enter 1 in vec2 wherever along the sequence $N_3N_4$ occurs. The location of a dinucleotide is defined as the location of the first base.

We then count the total number of times a '1' entry in vec1 lies $i$ bases before a 1 entry in vec2, or a 1 entry in vec2 lies $i$ bases before a 1 entry in vec1. To do so, we increment a variable $j$ from 1 to $50 - i - 1$, in steps of 1 and count the total number of times the boolean expression (vec1[j] = 1 AND vec2[j+i] = 1) OR (vec2[j] = 1 AND vec1[j+i] = 1) is TRUE. Say this number is N. N is the un-normalized value for $\rho(i)$ for the $N_1N_2$-$N_3N_4$ pair in the given sequence.

To normalize it, we generate a completely random set of 10,000 50 bp sequences and again calculate the unnormalized value of $\rho(i)$ for the $N_1N_2$-$N_3N_4$ pair among all these 10,000 sequences. We take the mean of these number among all 10,000 sequences and call the resulting mean M. The normalized value for $\rho(i)$ is then N/M.

#### Supplementary note 4:
#### Plotting details for figure 1e

We sort the 12,472 sequences in the random library according to increasing intrinsic cyclizability, and identify the 1,000 sequences with the lowest values of intrinsic cyclizability. We construct a matrix of dimension 1000 x 48. Each of the 1,000 rows corresponds to the 1,000 sequences, while within a row, the columns are populated by the values of $\rho(i)$ ($i$ = 1 to 48 bp) for the $N_1N_2$-$N_3N_4$ pair in that sequence. Plotted in blue is the column-wise average of this matrix. For each separation distance $i$, the blue curve thus represents $\langle\rho(i)\rangle$ averaged over the 1,000 sequences. The red curve is similarly obtained, except that we averaged over the 1,000 most cyclizable sequences in the random library.

#### Supplementary Note 5:
#### Details pertaining to figure 1f

For a given NN-NN pair, we calculated its helical separation extent as defined by equation (1) in the main text for all 12,472 sequences in the random library. We then fit the scatter plot of helical separation extent vs intrinsic cyclizability to a straight line and treat its slope as the contribution of the NN-NN pair towards intrinsic cyclizability. The slope values so obtained for all 136 NN-NN pairs is color coded and presented in figure 1f.

**Supplementary note 6:**
**Plotting details for figure 1g**

We first determined the sequences of the 4,904 2001 bp DNA fragments that straddle the +1 nucleosomal dyads (from position dyad – 2,000 till dyad + 2,000) of the identified 4,912 genes in *S. cerevisiae*. These sequences are listed in the supplementary file "Set_of_4912_Scerevisiae_sequences.txt".

Nucleosome occupancy is obtained from the re-analysis of earlier published data, as described before.

To obtain overall G/C content, we constructed a matrix of 4,912 rows and 2001 columns. Each row represents each of the 4,904 2,001 bp sequences. Each column is populated by 1 if at that location along the 2,001 bp sequence, a G or a C occurs. Otherwise, it is populated by 0. The plotted G/C content is the column-wise averaging of this matrix.

CpG and TpA contents are similarly obtained, except that the matrix has a dimension of 4,094 by 2,000, and each row is populated by 1 at the locations along the 2,001 bp sequence where the CpG or TpA dinucleotide occurs. See figure below for heatmaps of CpG and TpA contents (and nucleosome occupancy).

To plot the summed helical separation extent over the 10 NN-NN pairs that make the most negative contribution to intrinsic cyclizability ($\sum_{rigid} \sigma_{NN-NN}$), we first identified these pairs based on the values in figure 1f (and listed in "Contribution_of_HSE_to_IC_fig1.txt"). These pairs, in increasing order of contribution to intrinsic cyclizability, are: TT-GC, AA-GC, TA-GC, AT-GC, AA-CG, AA-CC, TT-GG, TT-CG, TT-CC, TA-CC. We tiled each of the 4,904 2,001 bp sequences as overlapping sets of 50 bp sequences, each offset from the neighbor by 1 bp. Therefore, the tiled sequences extend from positions 1 – 50, 2 – 51, 3 – 52, …, 1,952 – 2,001 along the original 2,001 bp sequence. We calculated the sum of the helical separation extents of these 10 NN-NN pairs for each of these tiled sequences, and assigned the value to the midpoint of the tile. Plotted is this value, averaged over all 4,904 genes, as a function of position from the +1 nucleosomal dyad.

Similarly, we obtained the mean summed helical separation extent over the 10 NN-NN pairs that make the most positive contribution to intrinsic cyclizability ($\sum_{flexible} \sigma_{NN-NN}$). The pairs were identified as before and are: CC-CC, GC-CG, GC-CC, TA-TT, AT-TT, AA-TA, AA-AA, AA-TT, TT-TT, GC-GC. The calculations were done as in the case of the negatively contributing pairs.

**Heatmaps of CpG and TpA contents, and nucleosome occupancy:**

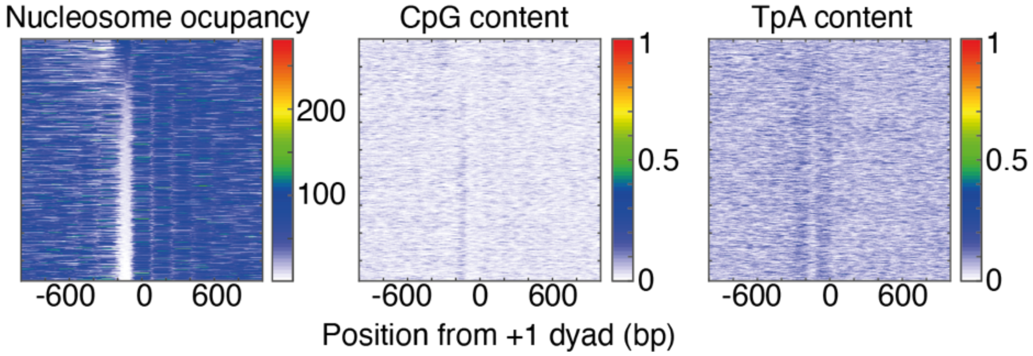


**Legend:** Heatmap of nucleosome occupancy, CpG content, and TpA content. Each row represents a gene. Genes are sorted according to increasing NDR strength (from top to bottom)

Sorting of genes was done as follows: For each gene, the mean nucleosome occupancy in the range [-161 bp, -101 bp] (0 being the dyad of the +1 nucleosome) was calculated. Genes were reverse-sorted according to this number, and considered to thus be sorted according to increasing NDR strength.

Obtaining CpG content for the plot above: For each gene, a row matrix containing 2000 columns was initialized. The 2,001 bp region straddling the +1 nucleosomal dyad of the gene was considered. The row matrix was populated with 1 at locations along this 2,001 bp region wherever ‘CG’ occurred. The row matrix was then smoothened with a rolling window average of 50 bp. Each row in the figure above corresponds to a color-coded version of this smoothened row matrix.

A similar procedure was followed to obtain the heatmaps of TpA content and nucleosome occupancy.

**Supplementary note 7:**
**Bendability quotients**

To obtain the bendability quotients for all 256 NNNN tetranucleotides, we closely followed the procedure outlined in supplementary note 2. As done in the case of dinucleotides (Fig. 1d, supplementary note 2), the 12,472 sequences were sorted and divided into 12 bins in order of increasing intrinsic cyclizability. The normalized number of times each tetranucleotide occurs in each of the bins was calculated and represented as a colormap in figure 2b.

For each of the 12 bins, we calculated the mean intrinsic cyclizability of the 1,039 sequences in the bins. Then, for each of the 256 tetranucleotides, we plotted its normalized content in each bin against the mean intrinsic cyclizability of the bin, fit it to a straight line, and defined the slope as the bendability quotient of that tetranucleotide.

#### Supplementary note 10:
#### One hot coding for DNA sequences

When DNA sequences were fed as inputs to a neural net, the sequence was represented numerically vis one-hot coding. We represented each nucleotide as a 4-element vector as follows:

A = [1,0,0,0]
T = [0,1,0,0]
G = [0,0,1,0]
C = [0,0,0,1]

Thus a 50 bp sequences is represented by 200 element vectors, each nucleotide in the sequence contributing 4 elements to the vector, in order. A neural net was trained to take as input such a 200-element vector and predict a single number: the intrinsic cyclizability (MATLAB). The net was trained using the tiling library dataset of measured intrinsic cyclizability values and used to predict the intrinsic cyclizabilities of sequences in the random library, the chrV library, and the cerevisiae nucleosomal library.

**Supplementary note 11:**
**The Linear Physical model**

In the first physical model we developed (Fig. 3b), intrinsic cyclizability was considered to be a linear combination of the number of times the 16 NN dinucleotides occur in the sequence:

$$C_0^i = A + \sum_{j=1}^{16} B_j N_j^i \qquad (2)$$

Here, $C_0^i$ is the intrinsic cyclizability of the $i^{th}$ sequence in the library, $N_j^i$ ($j$ = 1,…,16) is the content of the $j^{th}$ dinucleotide in the $i^{th}$ sequence, $B_j$ are regression coefficients, and A is a constant term. We subject it to the constraint $\sum_{j=1}^{16} N_j^i = 49$ for all $i$, since there are 49 possible dinucleotides in a 50 bp sequence.

To train the model and obtain the values of Bj and A, we considered the measured intrinsic cyclizability values of all sequences in the tiling library. For each value of $i$ (i.e., for each sequence in the library), we can write an equation of the form 2. As the number of sequences (~90,000) is much greater than the number of unknowns (the 16 $B_j$ values and $A$), the system is over-determined. We obtained the least square solution for $A$ and the 16 $B_j$s (MATLAB).

In the next model (Fig. 3c), we considered intrinsic cyclizability to be a linear combination of the 136 helical separation extents of the 136 NN-NN pairs:

$$C_0^i = A + \sum_{j=1}^{136} B_j \sigma_j^i \qquad (3)$$

. Here $C_0^i$ is the intrinsic cyclizability of the $i^{th}$ sequence in the library and $\sigma_j^i$ is the helical separation extent as defined by equation (1) of the $j^{th}$ NN-NN pair in the $i^{th}$ sequence of the library. $B_j$ are regression coefficients. As before, we performed multivariant linear regression to obtain the best fit values of $A$ and $B_j$ by using the measured intrinsic cycliability values of sequences in the tiling library as the training dataset.

Finally, we combined the two models into a single linear physical model (Fig. 3d) for intrinsic cyclizability as follows:

$$C_0^i = A + \sum_{j=1}^{16} B_j N_j^i + \sum_{j=1}^{136} C_j \sigma_j^i \qquad (4)$$

Here, where $B_j$ and $C_j$ are regression coefficients associated with the $j^{th}$ dinucleotide ($j$ = 1,…,16) and the $j^{th}$ NN-NN pair ($j$ = 1,…,136). As the equations are subject to the constraint $\sum_{j=1}^{16} N_j^i = 49$, We summed over only the first 15 $N_j$ values (those associated with all dinucleotides except CC), and consumed $N_{CC}$ (i.e., the 16$^{th}$ $N_j$) into the constant term. The final set of equations, therefore, had the form

$$C_0^i = A + \sum_{j=1}^{15} B_j N_j^i + \sum_{j=1}^{136} C_j \sigma_j^i \qquad (5)$$

For the purposes of figure 3d, we trained the model using the measured values of intrinsic cyclizability in the tiling library. However, for subsequent predictive purposes, we trained the model using the measured intrinsic cyclizability values of the random library and the methylated random library.

**Supplementary Note 13:**
**Non-linear physical model based on neural nets**

The neural net model where sequences are represented by one-hot coding does not provide any physical insight into how sequence features influence intrinsic cyclizability (supplementary note 10). We identified 152 sequence parameters that can be used to describe a 50 bp sequence (the 16 NN content terms and the 136 NN-NN helical separation extent perms) that influence its intrinsic cyclizability to various degrees. The linear physical model (supplementary note 11), however, assumed that intrinsic cyclizability of a 50 bp sequence is a linear combination of these 152 parameters (and a constant term). We now relax this condition of linearity. We still describe a sequence by this set of parameters, but then feed these parameters and input to a shallow neural net with a single hidden layer of 10 neurons to predict intrinsic cyclizability. The net is trained using measured intrinsic cyclizability values of the tiling library.

Because of the constraint that along any sequence, $\sum_{j=1}^{16} N_j = 49$ where $N_j$ is the NN content of the $j^{\text{th}}$ dinucleotide, only 15 of the 16 NN contents were fed as inputs (along with all 136 NN-NN helical separation extents).

#### Supplementary note 14:
#### Plotting details for figure 3f

The plot of measured intrinsic cyclizability as well as nucleosome occupancy averaged over the 576 genes, as a function of position from the dyad of the +1 nucleosome was obtained as described earlier.

The tiling library comprised 143 50 bp fragments for each of the 576 genes, that tile the region from 600 bp upstream till 400 bp downstream of the dyads of the +1 nucleosomes of these genes at 7 bp resolution (supplementary note 1). The intrinsic cyclizability values of these 82,368 fragments were predicted using the linear physical model trained using the random library (unmethylated) (supplementary note 11).

After predicting intrinsic cyclizability values of these fragments, we constructed a matrix of 576 rows and 143 columns. Each row represents a gene in the library, and its columns were populated with the predicted intrinsic cyclizability values of the 143 DNA fragments that tiled that particular gene. Plotted in grey is the column-wise average of this matrix. Plotted in red is the grey curve smoothened over a sliding window of 7 fragments. The 143 x-axis values represent the distances of the centers of the 143 fragments from the location of the +1 nucleosome dyad.

**Supplementary note 15:**
**Plotting details for figure 4a**

Various numbers of genes were identified in the case of the 4 organisms for which we had nucleosome occupancy data[8–10]. Genes were oriented in the direction of transcription and aligned according to the dyads of the +1 nucleosomes (in the case of *S. cerevisiae*[1] and *S. pombe*[11]) or to the TSSs (in the case of drosophila[9] and mouse[10]). For every gene, a 2001 bp region was considered, spanning the region (alignment point – 1,000) bp till (alignment point + 1,000) bp.

A process very similar to what is described for figure 3f was followed to obtain predicted intrinsic cyclizability (via the linear physical model trained using the random or the methylated random libraries) as a function of position, averaged over these genes for each organism. Briefly, a given 2,001 bp was tiled as a set of 50 bp fragments, each fragment offset from the neighbor by 7 bp. Intrinsic cyclizability values of these fragments were predicted using the linear physical model (and using parameters obtained by using the random library and the methylated random library for training), and the predicted values were aligned and averaged for each organism as done in figure 3f. G/C content was obtained as described for figure 1g.

#### Supplementary note 16:
#### Plotting details for figure 4b

We identified 19,900 CTCF binding sites in mouse[12] and identified a 2,020 bp region straddling the CTCF binding motif. The motif extends from position 1,000 till 1,019 along these 2,020 bp region. As the CTCF consensus motif is non palindromic, the sequences were all oriented in the same direction. As done in the case of figures 3f and 4a, we tiled all of these 2,020 bp sequences as a set of 50 bp sequences, each offset from its neighbor by 7 bp. We predicted intrinsic cyclizability values of these 50 bp sequences using the combined linear model (trained against the random library). The predicted intrinsic cyclizability values were aligned and averaged over the 19,900 sequences to obtain mean predicted intrinsic cyclizability as a function of position from the start of the CTCF consensus motif (top panel).

The top panel indicated that intrinsic cyclizability is not only high in the region defined by the CTCF consensus motif, but also for ~100 bp downstream. To verify that this is not an artifact of averaging, or of the fact that intrinsic cyclizability can only be defined for 50 bp fragments, we created an altered set of the 19,900 sequences. The sequences were identical in the 20 bp region of the CTCF consensus motif, but surrounding DNA was completely random in this set. We repeated the process described above and found that in this case, no extended region of higher intrinsic cyclizability than surrounding DNA is seen (Fig. 4b, bottom panel).

**Supplementary note 17:**
**Plotting details for figure 4c**

The Center Weighted Occupancy (CWO) in the 2,020 bp region straddling the 19,900 CTCF binding sites in mouse were obtained as reported earlier on the basis of chemical cleavage data[10]. For each site, the mean CWO value from position 1006-10 till 1006+10 was calculated and considered to be the "CTCF nucleosome occupancy" of the CTCF binding site (the CTCF consensus motif extends from position 1,000 till 1,019). From among the 19,900 sites, the 1,000 sites that had the lowest and highest values of "CTCF nucleosome occupancy" were identified. Plotted is the mean CWO as a function of position from the CTCF consensus motif averages over these two sets of CTCF sites. Also plotted is the mean CWO as a function of position averaged over all 19,900 sites.

#### Supplementary Note 19:
#### Plotting details for figure 5e

The 13,300 200 bp DNA sequences around the dyads of gene body nucleosomes identified in figure 5b were written one below the other, creating a matrix of 13,300 rows and 200 columns. Because the sequences lie entirely within coding regions, they could all be translated in-frame. Each sequence was replaced by the equivalent amino-acid sequence, and all three nucleotides of a codon were replaced by the same amino acid it codes for. For example, the codon TAT codes for Y. Thus if TAT is an in-frame codon that appears along a row in the matix, it was replaced by YYY. Each amino acid was subsequently replaced by its slope as represented in figure 5d and listed in supplementary note 18. Plotted in figure 5e is the column wide mean and s.e.m. of this resulting matrix.

**Supplementary Note 20:**

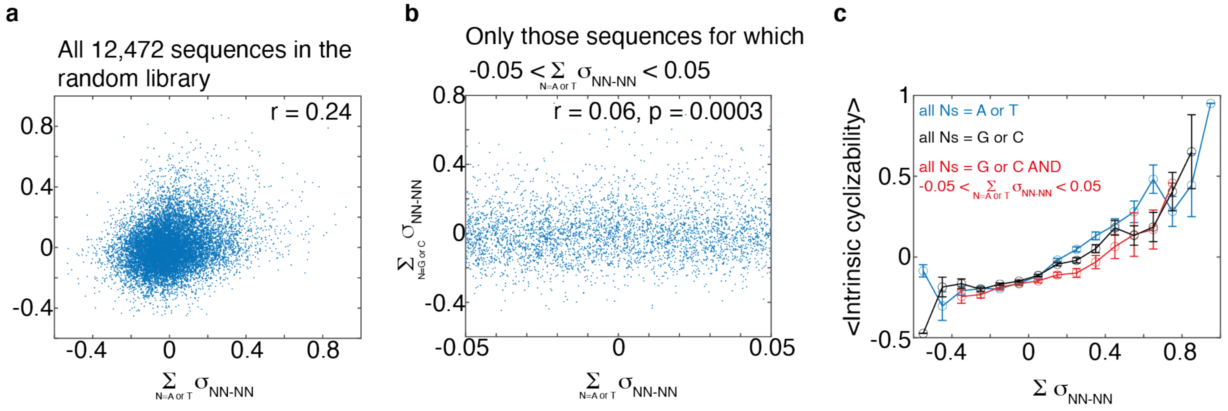


In figure 1f, we see that NN-NN pairs for all Ns = A or T tent to make a positive contribution to intrinsic cyclizability, in accordance with earlier results that polyA tracts repeated at the helical repeat yield curved molecules and repeated at half the helical repeat yield straight molecules.

Figure 1f also shows that NN-NN pair for all Ns = G or C also tend to make a positive contribution to intrinsic cyclizability. We wanted to understand if this is indeed a new result, or it is it just because seuqneces which have a high helical separation extent for A/T containing NN-NN pairs also tend to have a high helical separation extent for G/C containing NN-NN pairs.

For every sequence in the random library, we calculated the sum of the helical separation extents of all NN-NN pairs for N = A or T. Likewise, we also calculated this sum for all N = G or C pairs. As expected, there is a weak by significant correlation between the two among all sequences in the random library (panel a above). This really just reflects the fact that if a sequence has high A/T content every 10 bp, it is likely, by exclusion, to have high G/C content also every 10 bp, offset in phase by 5 bp.

When sequences were binned and sorted according to increasing summed helical separation extents over A/T containing as well as G/C containing NN-NN pairs, we find that in either of these cases, increasing binned helical separation extent leads to increased intrinsic cyclizability (blue and black curves in panel c above). This again is identical to the result depicted in figure 1f that A/T or G/C containing NN-NN pairs make significantly positive contributions to intrinsic cyclizability.

We now decided to select a subset of sequences for which the summed helical separation extent of A/T containing NN-NN pairs has very little variation (it lies between -0.05 and 0.05). Among these sequences, there is very little correlation between summed helical separation extents of the A/T containing vs the G/C containing NN-NN pairs (panel b above). However, even among these sequences which all have very similar values of the summer helical separation extent of A/T containing pairs, increasing summed helical separation extents of G/C containing pairs leads to increased intrinsic cyclizability (panel c above, red curve).

These findings suggest that positive correlation between helical separation extents of G/C containing NN-NN pairs and intrinsic cyclizability is not just because helical separation extents of G/C containing and A/T containing pairs are among themselves correlated.

### Supplementary Note 21:

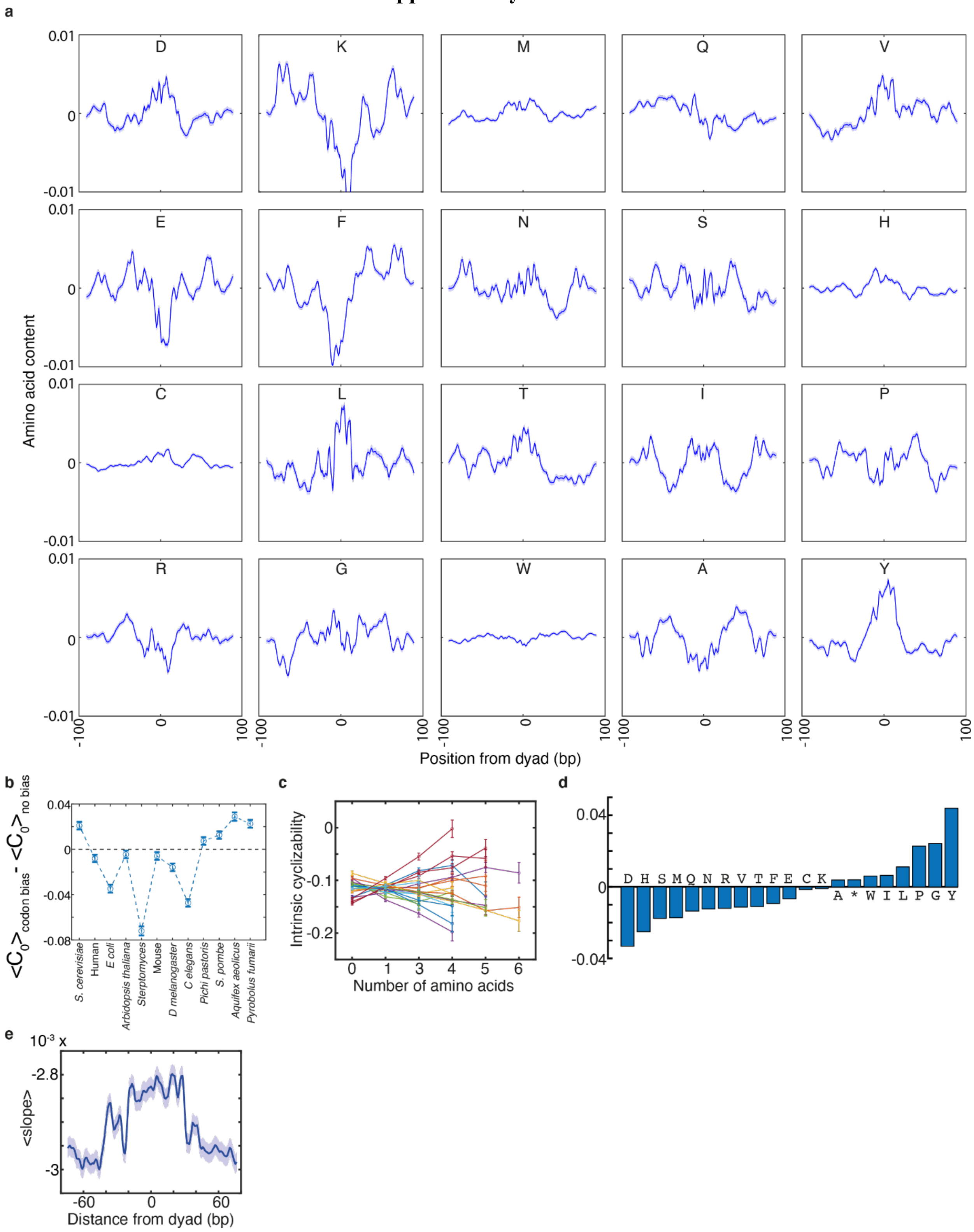

**Panel a:** contents of all 20 amino acids along the 13,310 gene-body nucleosomes identified in the context of figure 5b.

**Panel b:** Figure 5b indicates that the effect of codon-bias in *S. cerevisiae* is the overall increase the intrinsic cyclizability of DNA. We asked if this result is generally true across organisms. To do so, we created a set of 1,000 random 17 amino acid long polypeptide sequences. We converted these 1,000 amino acid sequences to 1,000 DNA sequences by randomly selecting synonymous codons. We did this process 10 times, generating a total of 10 sets of 1,000 DNA sequences each. For the first 5 sets, random selection between synonymous codons were biased by the natural codon usage frequency of a particular organism, while in the subsequent 5 sets, all synonymous codons were selected with equal probability. We predicted the intrinsic cyclizabilities of every sequence in all the 10 sets. For sequence 1, we tabulated the following 5 numbers:

[Intrinsic cyclizability of sequence1 in set 1 – Intrinsic cyclizability of sequence 1 in set 6; Intrinsic cyclizability of sequenc 1 in set 2 – Intrinsic cyclizability of sequence 1 in set 7; …; Intrinsic cyclizability of sequence 1 in set 5 – Intrinsic cyclizability of sequence 1 in set 10].

We note the mean of this set of numbers, and accordingly repeat it for all 1,000 sequences. This process was repeated for various organisms, each organism being characterized by a different codon useage frequency. Plotted in panel b is the mean of these set of 1,000 numbers and the associated s.e.m., for various organisms.

We find that codon usage bias does not necessarily result in increased intrinsic cyclizability over random codon assignment, however, the difference is significantly different from 0. We were unable to correlate the direction of this bias with any other feature describing the organism.

Panels c-e: In figures 5d, we tabulated the slope associated with an amino acid (i.e., the measure of its contribution towards intrinsic cyclizability) based on fitting to a straight line the intrinsic cyclizability of 50 bp DNA sequences in coding regions of S. cerevisiae chromosome V vs the number of times the amino acid in question occurs in its equivalent in-frame translated amino acid sequences contain. We then find that amino acids are not randomly distributed along nucleosomes: the mean slope of amino acids along gene body nucleosomes is high near the dyad and low near the edges.

An even stronger claim that the requirement for DNA to be highly flexible near nucleosome dyads constrains the possible amino acid sequences can be made if the slope values associated with the amino acids can be obtained not from the coding region of *S. cerevisiae* genome, but from totally random DNA. We therefore considered the 12,472 sequences in the random library, and translated each sequence in all three possible frames. To all these amino acid sequences, we assigned the same value of measured intrinsic cyclizability as was measured for the original DNA sequence. We therefore obtained a total of 37,416 amino acid sequences as their associated intrinsic cyclizability values. We repeated the analysis as done in figure 5d, and plotted separately for each amino acid, the mean intrinsic cyclizability as a function of the number of times the amino acid occurs in the sequences (panel c above), and tabulated the slope for every amino acid (panel d above). Then, as done in figure 5e, we plotted the mean slope of amino acids as a function of position along gene body nucleosomes, averaged over all identified 13,310 nucleosomes that are classified as +5 – +9 in *S. cerevisiae*.

**Supplementary Note 22:**
**G/C content cannot explain the nucleosome organization around CTCF binding sites**

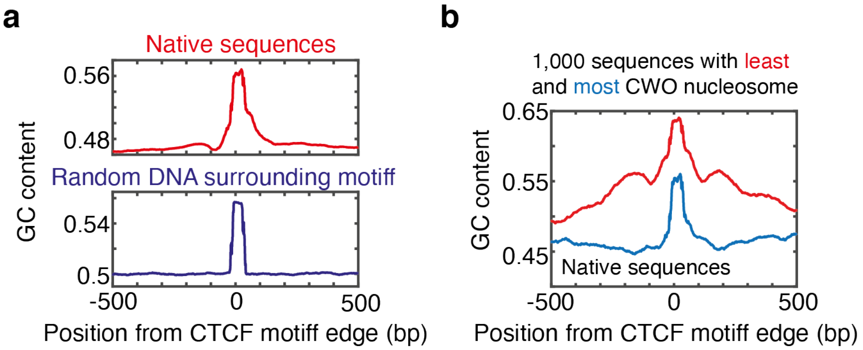


Panel a: Same as Fig. 4b, except that average G/C content, instead of predicted intrinsic cyclizability, is being plotted as a function of position from CTCF binding sites.

Panel b: Same as Fig. 4d, except that G/C content, instead of predicted intrinsic cyclizability, is plotted as a function of position from the CTCF binding sites, averaged over the 1,000 sites that have the least (blue) or most (red) Center Weighted Occupancy (CWO) value of the nucleosome that occupies the CTCF site.

Panel a by itself would suggest that high G/C content might be indicative of high nucleosome density because the CTCF binding site is known to harbor a well-positioned nucleosome. However, this would be inconsistent with panel b, where CTCF sites with the least CWO nucleosomes seem to have even higher G/C content than sites with the most CWO nucleosomes. In contrast, in Fig. 4d, we see that CTCF sites with the least CWO nucleosomes have lower intrinsic cyclizability than CTCF binding sites with the highest CWO nucleosomes, consistet with Fig. 4a, where we find that CTCF sites, on average, have higher intrinsic cyclizability than neighboring regions. Further, the heightened contrast between intrinsic cyclizability at the CTCF site and in the immediate vicinity seen for those sites that are occupied by high CWO nucleosomes, indicates a mechanical basis for why these nucleosomes have high CWO scores and are well positioned. No such difference in G/C content contrast is seen (panel b).

Further, from Fig. 4a, we find that in the case of S. pombe, nucleosome dyads are clearly marked by regions of log CpG content, which is again inconsistent with panel a above, given a nucleosome is known to reside at the CTCF binding site.

#### Supplementary Note 23:
#### Plotting details for Fig. 1a

The random library as well as the methylated random library had 7 “control” sequences introduced which were fully unmethylated in both cases. As intrinsic cyclizability is defined up to an additive constant, we used these sequences to adjust the cyclizability values of sequences in the methylated random library, to allow for comparison with sequences in the random library.
